# Spatial proteomics identifies altered thick ascending limb cells as drivers of kidney disease through uromodulin retention

**DOI:** 10.64898/2026.10.09.756985

**Authors:** Azuma Nanamatsu, George J. Rhodes, William S. Bowen, Angela R. Sabo, Shehnaz Khan, Mahla Asghari, Dillen L. Wischmeier, Daria Barwinska, Takashi Kudo, Azusa Tomioka, Hiroaki Sakaue, Atsushi Kuno, Ana C. Ricardo, James P. Lash, Katherine R. Tuttle, Pierre C. Dagher, Jonathan Himmelfarb, Kaice A. LaFavers, Luca Rampoldi, Tarek M. El-Achkar

**Author notes:** Corresponding authors: Azuma Nanamatsu, Division of Nephrology and Hypertension, Indiana University School of Medicine 950 W. Walnut, R2 202,Indianapolis, IN 46202., Tarek M. El-Achkar, Division of Nephrology and Hypertension, Indiana University School of Medicine 950 W. Walnut, R2 202, Indianapolis, IN 46202.

## Abstract

Chronic kidney disease (CKD) is increasingly prevalent and is a leading cause of mortality. Recent transcriptomic studies have identified the persistence of injury-induced renal tubular epithelial cell states, yet the cell populations that drive CKD remain unresolved. Uromodulin (UMOD) is the most abundant urinary secreted protein expressed exclusively in thick ascending limb (TAL) epithelial cells in the kidney. UMOD is strongly associated with CKD risk in genome-wide association studies, suggesting a broader, yet unexplored role of UMOD and TAL cells in CKD pathogenesis. Here, using spatial proteomics, we identified an injury-induced TAL cell state marked by intracellular UMOD accumulation. Spatially resolved multimodal integration revealed that UMOD accumulation occurred without transcriptional upregulation. Impaired urinary UMOD secretion further supported pathological intracellular retention. These altered TAL cells with UMOD retention exhibited inflammatory signaling, formed profibrotic niches enriched with M2-like macrophages across renal compartments, and were associated with CKD progression in multiple experimental models. Reducing the burden of UMOD retention diminished altered TAL cells and their profibrotic microenvironments, and attenuated CKD development. A similar altered TAL state was identified in human kidney disease and was associated with worse kidney function. Together, we identify pathological UMOD retention and altered TAL cells as key drivers of CKD development and potential therapeutic targets. This study also highlights that protein-level alterations can redefine disease-associated cell states beyond transcript profiling.

## INTRODUCTION

The kidney is a highly organized organ composed of a cortex and a multi-compartment medulla (**Fig. 1A**). This intricate spatial organization enables diverse functions that maintain homeostasis, such as glomerular filtration and urine concentration. Chronic kidney disease (CKD), defined by persistent abnormalities in kidney function or structure, is increasingly prevalent, affecting approximately 800 million people worldwide. CKD markedly increases risks of end-stage renal failure, cardiovascular disease and premature mortality(*1*, *2*). Current CKD therapies include renin-angiotensin system (RAS) inhibitors(*3*), sodium-glucose cotransporter-2 (SGLT2) inhibitors(*4*), non-steroidal mineralocorticoid receptor antagonists(*5*), and glucagon-like peptide-1 receptor agonists(*6*). These drugs ameliorate kidney damage by targeting shared final common pathways, such as hemodynamic and metabolic disturbances, inflammation, and fibrosis, irrespective of inciting injuries. Despite these important therapeutic advances, substantial residual risks of kidney failure, major adverse cardiovascular events and mortality remain, underscoring the need to better define the mechanisms driving CKD development and progression. Acute kidney injury (AKI), caused by ischemia, nephrotoxins, or sepsis, is increasingly recognized as a key driver of CKD through a maladaptive repair process known as AKI-to-CKD transition(*7*). However, the underlying mechanisms remain incompletely understood. Defining these mechanisms will be essential for developing therapeutic strategies to prevent CKD development or halt its progression.

**Fig. 1.**
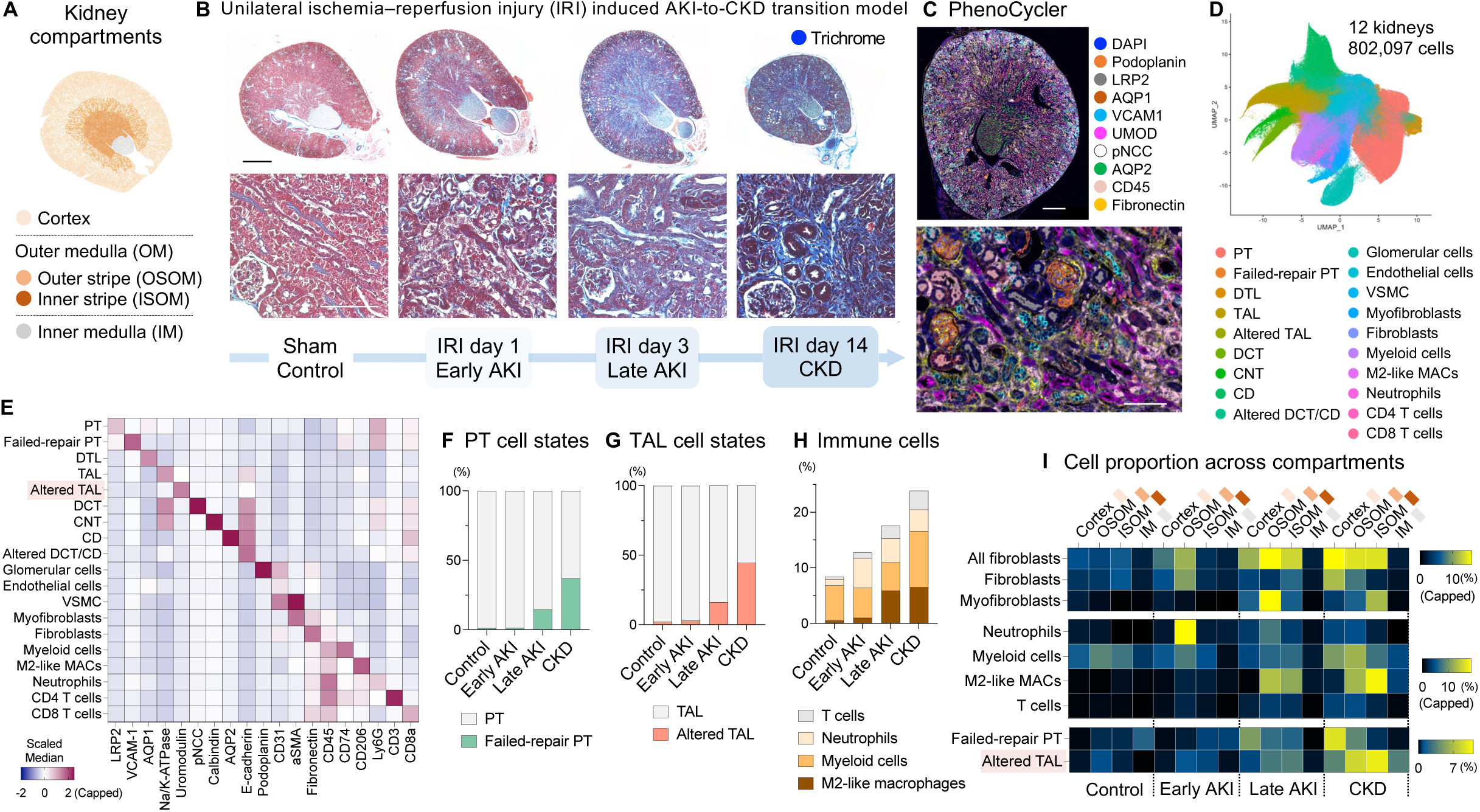
Spatiotemporal cellular landscape during AKI-to-CKD transition revealed by spatial proteomics. Wild-type mice underwent sham surgery (control) or unilateral ischemia–reperfusion injury (IRI) with 23-minute clamp time and were harvested at day 1, day 3 and day 14. n = 3 per condition. (A) Visualization of kidney compartments. (B) Masson’s trichrome staining of mouse kidneys. Bars = 1 mm or 100 μm (zoom in). (C-I) PhenoCycler protein imaging was performed on control, early AKI, late AKI and CKD kidneys (n = 3 per group). (C) Representative PhenoCycler image. Bar = 500 μm or 100 μm (zoom in). (D) UMAP plot showing cell types. (E) Heatmap displaying median expression of marker proteins across clusters. (F) Proportion of failed-repair PT cells among PT cells at each disease stage. (G) Proportion of altered TAL cells among TAL cells at each disease stage. (H) Proportion of immune cell subtypes at each disease stage. (I) Heatmap showing proportion of each cell type across disease stages and anatomical regions.

Recent single-cell and spatial transcriptomics studies have revealed injury-induced tubular epithelial cell states in both humans and mice(*8–14*). These cell populations are proposed to drive CKD progression by forming pathological niches and activating immune cells and fibroblasts. Among tubular segments, maladaptive states are primarily observed in proximal tubule (PT) and thick ascending limb (TAL) cells, the two most abundant epithelial cell types in the kidney. While studies of CKD mechanisms have largely focused on PT cells which are highly vulnerable to acute injury, recent human studies have highlighted the association between maladaptive TAL states and reduced kidney function(*9*, *15*). However, the molecular mechanisms governing altered TAL states and their causal roles in CKD development remain unresolved.

Importantly, these disease-associated cell states have been defined largely by transcriptomic profiling. However, transcript abundance does not necessarily reflect protein localization or intracellular processing, which are the key determinants of cellular function and disease mechanisms. Whether spatial proteomics can redefine disease-associated cell states in kidney disease remains largely unexplored.

Uromodulin (UMOD), a TAL-specific secretory protein with strong genetic links to CKD, has emerged as a potential molecule implicated in CKD pathogenesis. UMOD is the most abundant protein secreted into the urine(*16*). Numerous genome-wide association studies (GWASs) have consistently shown that common single-nucleotide polymorphisms (SNPs) in the non-coding regions of the *UMOD* gene, driving high UMOD expression(*17*), are associated with an increased risk of CKD incidence and progression(*18–21*). Because this association is highly robust and reproducible, these *UMOD* variants are often used as benchmarks in CKD GWASs. Another important aspect of UMOD is that rare pathogenic mutations in the coding regions cause a monogenic form of CKD, known as autosomal dominant tubulointerstitial kidney disease-UMOD (ADTKD-*UMOD*)(*22*). In ADTKD-UMOD, mutant UMOD misfolds and accumulates intracellularly, inducing ER stress, proinflammatory and profibrotic signaling that drive CKD development. Whether increased UMOD abundance and aberrant cellular processing contribute to the common AKI-to-CKD transition remains unknown.

## RESULTS

### Spatiotemporal cellular landscape during AKI-to-CKD transition revealed by spatial proteomics

We first aimed to generate a longitudinal spatial proteomic atlas of AKI-to-CKD transition to delineate the temporal changes in the underlying cellular architecture. We used a well-established murine unilateral ischemia–reperfusion injury (IRI) model(*23–25*) and harvested at multiple time points (**fig. S1A**). In this model, acute necrosis was observed at day 1, consistent with the established peak of initial injury(*26*), followed by collagen deposition at day 3. By day 14, global fibrosis and renal atrophy, as reflected by kidney weight loss, were evident (**Fig. 1B** and **fig. S1B**), representing cardinal features of AKI-to-CKD progression(*23*). Fibrosis was prominent in the cortex and outer medulla (OM), including outer stripe of the outer medulla (OSOM) and inner stripe of the outer medulla (ISOM) (**Fig. 1, A and B**). In this study, we defined IRI day 1, day 3, and day 14 as early AKI, late AKI, and CKD, respectively.

We performed PhenoCycler spatial protein imaging on formalin-fixed paraffin-embedded (FFPE) whole kidney tissue sections (n = 3 per condition) (**Fig. 1C**) using the antibody panel shown in **table S1**. We analyzed 802,097 cells from 12 kidneys and graph-based clustering identified 19 cell types (**Fig. 1, D and E** and **fig. S1, C and D**). Cell composition analysis showed that the AKI-to-CKD transition was marked by progressive accumulation of immune cells and fibroblasts (**fig. S1E**).

Among altered tubular subtypes, we observed VCAM-1-positive failed-repair PT (FR-PT) cells during disease progression (**Fig. 1, E and F**), consistent with previous reports using single-nucleus and spatial transcriptomics on murine IRI kidneys(*8*, *14*). We also identified altered TAL (aTAL) cells which emerged at late AKI stage and expanded with CKD progression (**Fig. 1G**). aTAL cells showed reduced Na/K-ATPase and E-cadherin expression compared to healthy TAL cells (**Fig. 1E**), consistent with a partial dedifferentiated state. Notably, these cells were marked by elevated UMOD protein levels, a feature not previously resolved by transcriptomic analyses.

In this dataset, immune cells were classified into five cell types (myeloid cells, M2-like macrophages, neutrophils, CD4 T cells, and CD8 T cells) based on the expression of specific markers. Due to the lack of dendritic cell markers, the “myeloid cells” population was presumed to include dendritic cells and non–M2-like macrophages. Among immune cells, M2-like macrophages emerged during late AKI (**Fig. 1H**), consistent with pro-resolution subtypes during the repair phase(*24*). However, their persistence into the CKD phase beyond the expected repair period (**Fig. 1H**) strongly suggests a maladaptive transition to profibrotic subtypes(*27*). During late AKI and CKD, a subset of fibroblasts exhibited a myofibroblast phenotype, a key effector of fibrosis(*28*) (**Fig. S1F**).

Compartment-resolved cell composition analysis showed that fibroblasts predominantly accumulated across cortex, OSOM, and ISOM during CKD development (**Fig. 1I**), consistent with histological changes (**Fig. 1B**). In particular, myofibroblasts were preferentially localized to the ISOM in CKD. M2-like macrophages predominantly accumulated in the OSOM and ISOM. These data highlight the importance of the two OM regions, alongside the cortex, in CKD development. Regarding altered tubular states, failed-repair PT cells were enriched in the cortex, consistent with recent spatial transcriptomic observations(*14*, *15*). In contrast, aTAL cells were detected in the cortex but were more abundant in the OSOM and ISOM. The distribution of aTAL cells is similar to that of M2-like macrophages, suggesting potential interactions between aTAL cells and profibrotic immune cells during CKD progression.

### Altered TAL cells exhibit increased UMOD protein without transcriptional upregulation

To define the transcriptomic signature of aTAL cells identified by PhenoCycler protein imaging, we leveraged single-cell-resolution Visium HD spatial transcriptomics as a bridge between spatial proteomics and single-nucleus RNA sequencing (**Fig. 2A**). Visium HD spatial transcriptomics was performed on serial sections from one control kidney and one CKD kidney (IRI day 14) that were analyzed by PhenoCycler. We analyzed 209,274 cells from two kidneys with an average of 1,274 total transcripts and 926 individual genes detected per cell. Cell clustering identified 23 cell types with distinct gene expression profiles and spatial localization (**fig. S2, A to D**). For TAL cells, three TAL subtypes were identified: two regionally distinct healthy populations (TAL #1 and TAL #2) and a CKD-specific altered TAL population (**fig. S2, D and E**).

**Fig. 2.**
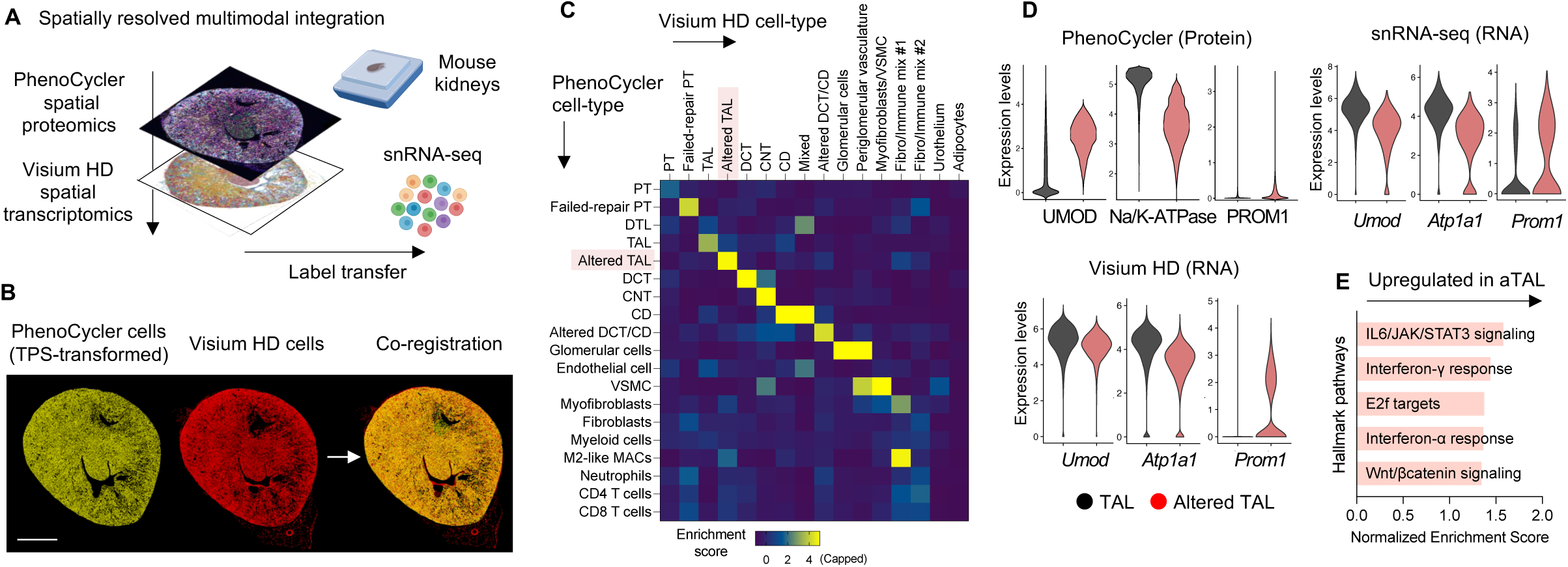
Altered TAL cells exhibit increased UMOD protein without transcriptional upregulation. (A) PhenoCycler spatial protein imaging and single-nucleus transcriptomic data were integrated using single-cell scale Visium HD spatial transcriptomics as an anchor. (B) Spatial registration of PhenoCycler (light yellow) and Visium HD (red) cells using landmark-based thin plate spline (TPS) transformation. Bar = 1mm. (C) Comparison of cell annotations between PhenoCycler cells and their spatially corresponding Visium HD cells. To calculate the enrichment score, the proportion of Visium HD annotations assigned to each PhenoCycler cell was subtracted by the overall frequency of each cell type in the Visium HD sample, and then divided by this overall frequency. (D) Expression of key markers in TAL and altered TAL cells measured by PhenoCycler (log1p-transformed), Visium HD, and snRNA-seq. (E) Gene Set Enrichment Analysis (GSEA) for genes upregulated in altered TAL cells compared with (healthy) TAL cells.

To determine whether altered TAL cells defined by PhenoCycler protein imaging correspond to those identified by Visium HD spatial transcriptomics, we performed spatial registration to align the cell types defined by the two modalities. After thin-plate spline–based registration (**Fig. 2B**), each PhenoCycler cell was assigned the nearest-neighbor Visium HD cell. 92.9% of PhenoCycler cells had a corresponding Visium HD cell within 10 μm, supporting the fidelity of the spatial registration (**fig. S2F**). We then compared cell annotations between spatially corresponding PhenoCycler and Visium HD cells. Major cell types showed overall consistency between the two modalities, demonstrating the accuracy of both cell annotation and spatial registration (**Fig. 2C**). Comparison of TAL subtypes also showed marked concordance across modalities, suggesting that aTAL cells identified by PhenoCycler correspond to those identified by Visium HD. This was further supported by similar spatial distributions of aTAL cells across the two platforms (**fig. S2G**). Thus, the gene expression signature of Visium HD aTAL cells likely corresponds to aTAL cells identified by PhenoCycler.

To gain even higher transcriptome coverage, we next conducted single-nucleus RNA sequencing (snRNA-seq). The FFPE-compatible GEM-X Flex Gene Expression platform enabled snRNA-seq from the same tissue blocks used for PhenoCycler and Visium HD analysis. After filtering, the snRNA-seq dataset included 30,962 cells from 6 kidneys, with an average of 3,071 total transcripts and 1,358 individual genes detected per cell. Cell clustering identified 22 cell types, including two TAL populations that may reflect regional variation rather than cell states (**fig. S3, A to C**). We then transferred TAL subtype signatures from the Visium HD dataset onto TAL populations in the snRNA-seq dataset. Label transfer achieved high confidence, with 94.25% of cells assigned prediction scores ≥ 0.7 (**fig. S3D**). This integrated approach enabled reliable identification of aTAL cells in the snRNA-seq dataset, which were also enriched in CKD (**fig. S3, E and F**).

Head-to-head comparison of mRNA and protein expression signatures between aTAL and healthy TAL cells across three modalities (PhenoCycler, Visium HD, and snRNA-seq) revealed a consistent reduction in Na/K-ATPase (*Atp1a1*) in aTAL cells (**Fig. 2D**). Similarly, PROM1 (*Prom1*), a dedifferentiation marker and a key marker of adaptive/maladaptive TAL cells in humans(*9*) was consistently elevated in aTAL cells. By contrast, UMOD showed a marked increase in protein without transcriptional upregulation, indicating post-transcriptional dysregulation of UMOD in aTAL cells.

We next conducted differential gene expression analysis between aTAL and healthy TAL cells using snRNA-seq dataset (**fig. S3G**; differentially expressed genes are listed in **Data S1 and S2**). The top up-regulated gene in aTAL cells was *Dcdc2a*, another key marker of adaptive/maladaptive TAL cells in humans(*9*). aTAL cells also showed increased expression of *Lcn2* (NGAL), an ER stress-responsive gene(*29*) that is induced by the intracellular accumulation of ADTKD-UMOD-causing mutant UMOD(*30*, *31*). In contrast, the most downregulated gene in aTAL cells was *Egf,* which encodes epidermal growth factor (EGF), a secreted protein predominantly expressed in TAL cells within the kidney(*8*, *32*). Urinary EGF has emerged as a promising biomarker for CKD in humans(*33*, *34*). Pathway analysis showed that aTAL cells possess an inflammatory transcriptomic signature (**Fig. 2E**).

Collectively, these data suggest that high UMOD protein levels in aTAL cells are associated with aberrant cellular processing rather than enhanced transcription. In addition, aTAL cells exhibited reduced Na/K-ATPase and increased PROM1 expression at both the protein and transcript levels, together with increased *Lcn2*, decreased *Egf*, and inflammatory gene expression signatures.

### Altered TAL cells exhibit intracellular UMOD accumulation and impaired urinary secretion

We next investigated intracellular UMOD protein processing in aTAL cells. We first examined the distribution of TAL and aTAL cells and corresponding UMOD protein expression in the IRI trajectory PhenoCycler dataset (**Fig. 3A**). In control mice, (healthy) TAL cells are the dominant state of TAL cells, and UMOD staining was predominantly localized to the apical membrane, consistent with previous observations(*35–38*). In contrast, aTAL cells emerged during the AKI-to-CKD transition and were associated with intracellular UMOD accumulation (**Fig. 1G and 3A**). Immunoblot analysis of whole kidney lysates confirmed that UMOD protein levels were significantly increased during the AKI-to-CKD transition (**Fig. 3B**), consistent with PhenoCycler imaging data. qPCR of whole kidney lysates showed that *Umod* mRNA levels were not upregulated (**fig. S4A**), in line with the observation in aTAL cells in Visium HD and snRNA-seq analyses. We have recently reported an alternatively spliced intracellular isoform of UMOD, which is induced in the early phase of mild AKI(*39*). This variant was not induced in this CKD model driven by severe AKI (**fig. S4B**), consistent with our report(*39*), thereby excluding the possibility that enhanced splicing increased intracellular protein levels. Confocal immunofluorescence images showed diffuse intracellular accumulation of UMOD in CKD, in contrast to its apical localization observed in controls (**fig. S4C**), consistent with the PhenoCycler imaging data (**Fig. 3A**). Note that NKCC2, a TAL-specific transporter, maintained its apical localization in CKD (**fig. S4C**), suggesting a unique impairment of UMOD trafficking, rather than a global disruption of epithelial polarity or membrane trafficking.

**Fig. 3.**
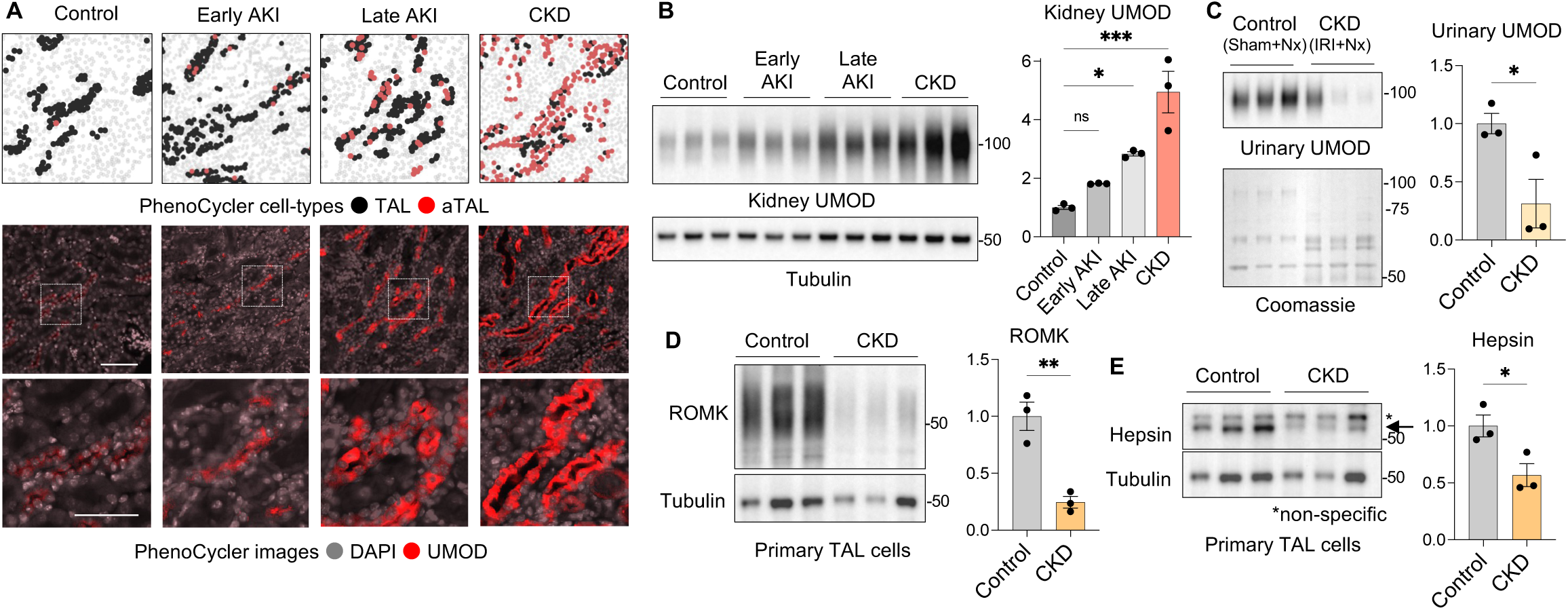
Altered TAL cells exhibit intracellular UMOD accumulation and impaired urinary secretion. (A) Spatial distribution of TAL subtypes and corresponding UMOD staining of PhenoCycler dataset on IRI kidneys. Bars = 100 μm or 50 μm (zoom in). (B) Immunoblotting of kidney UMOD in IRI kidneys. n = 3 per group. (C) A modified IRI protocol including contralateral nephrectomy 24 hours before harvest was applied to assess urinary UMOD secretion from IRI kidneys. Immunoblotting showing urinary UMOD in control and CKD mice. Urine was applied according to urinary creatinine concentration for normalization. n = 3 per group. (D,E) Primary TAL cells were isolated from control and CKD mouse kidneys. (D) Immunoblotting of ROMK in TAL cells in CKD. n = 3 per group. (E) Immunoblotting of Hepsin in TAL cells in CKD. n = 3 per group. Values are represented as mean ± SEM. *p < 0.05, **p < 0.01, ***p < 0.001.

We next sought to assess urinary UMOD secretion from the IRI-CKD kidney. Since urine collected from unilateral IRI mice is derived from both the injured and contralateral kidneys, we performed contralateral nephrectomy 24 hours before harvest (**fig. S4D**), an established approach for functional assessment of the injured kidney(*23*, *40*). This method confirmed declined renal function in the IRI-induced CKD model (**fig. S4E**). UMOD protein was accumulated in the kidney (**fig. S4F**), similar to the IRI model without nephrectomy. In this setting, urinary UMOD was significantly reduced in CKD compared to controls, both with and without normalization to urinary creatinine (**Fig. 3C** and **fig. S4G**). These data suggest that intracellular UMOD retention is caused by impaired extracellular secretion.

To investigate the mechanisms underlying disrupted UMOD secretion in CKD, we examined two molecules indispensable for urinary UMOD secretion: ROMK(*37*), a K⁺ channel essential for maintaining transepithelial voltage in TAL cells, and hepsin(*35*, *36*), a membrane-bound serine protease. Knockout mice lacking these genes exhibit impaired urinary secretion and intracellular accumulation of UMOD(*35–37*) and are associated with ER stress and tubular damage signatures(*36*). To this end, we examined primary TAL cells from control and CKD kidneys, as their molecular signatures in CKD are expected to predominantly reflect those of aTAL cells due to their enrichment (**Fig. 1G, fig. S2E** and **S3F**). UMOD protein accumulation was observed in CKD-TAL cells (**fig. S5A**), in line with bulk kidney analysis. In this setting, both ROMK and hepsin protein expression were reduced in CKD-TAL cells (**Fig. 3, D and E**). Corresponding changes were observed in the mRNA expression of *Kcnj1* (ROMK) and *Hpn* (Hepsin) (**fig. S5B**). Visium HD and snRNA-seq datasets confirmed decreased expression of these genes in aTAL cells (**fig. S5, C and D**). These data suggest that UMOD retention in aTAL cells during CKD may be explained, in part, by the combined reduction of key components of the UMOD secretory machinery. High-resolution confocal imaging revealed UMOD accumulation in both the ER and Golgi apparatus (**fig. S5E**). Similar intracellular accumulation patterns have been reported in *Romk*- and *Hpn*-deficient mice(*36*, *37*), raising the possibility that defective secretion induces UMOD accumulation along the secretory pathway in CKD. This contrasts with ADTKD-UMOD, where misfolded UMOD predominantly accumulates in the ER.

CKD-TAL cells exhibited increased protein levels of spliced XBP1 and ATF4 (**fig. S5F**), suggesting enhanced ER stress, which may provide a link between UMOD retention and the inflammatory signature in aTAL cells. These data suggest that CKD may share some pathological features with ADTKD, including intracellular UMOD accumulation, cell stress, and inflammatory signatures, despite distinct mechanisms of UMOD retention.

To examine the broader relevance and generalizability of intracellular UMOD retention in CKD, we studied additional experimental models. First, we used aristolochic acid nephropathy (AAN) model (**fig. S6A**), a clinically relevant nephrotoxin-induced CKD model that closely mimics human disease(*41*). Our previous work showed that both *Umod* and spliced *Umod* mRNA levels were unchanged in this model(*39*). We have now further characterized this model and confirmed signatures associated with CKD (**fig. S6, B to D**). Similar to IRI, AAN model exhibited intracellular UMOD accumulation with decreased urinary UMOD secretion (**fig. S6, E to G**). To confirm the presence of aTAL cell state in AAN, we performed PhenoCycler protein imaging on AAN kidneys (**fig. S6, H to N** and **table S2**). Uninjured kidneys were included as controls, and newly collected IRI day 14 kidneys were used as benchmarks. We identified 19 cell types, similar to those observed in the IRI trajectory dataset. Similar to IRI kidneys, AAN kidneys exhibited increased fibroblasts and immune cells, particularly M2-like macrophages (**fig. S6, K and L**). aTAL population was expanded in both IRI and AAN kidneys (**fig. S6N**).

We next examined 5/6 nephrectomy model (**fig. S7, A to E**), which recapitulates progressive CKD driven by compensatory adaptation of the remaining nephrons after nephron loss(*42*). Intracellular UMOD retention was also observed in 5/6 nephrectomy model (**fig. S7, D and E**) in line with IRI and AAN kidneys. Cumulatively, our findings support that expansion of aTAL cells with intracellular UMOD accumulation and impaired urinary secretion is a common feature of CKD models.

### Altered TAL cells form CKD-specific niches with M2-like macrophages

We next investigated the aTAL microenvironment during the AKI-to-CKD transition to link local cell states with kidney-wide disease progression. We performed cellular neighborhood (niche) analysis using the IRI trajectory PhenoCycler dataset. Neighboring cells were defined using a 30 μm radius (∼2 cell layers), corresponding to the expected range of juxtacrine and paracrine interactions. This parameter was chosen based on a recent seqFISH spatial transcriptomic study of IRI mouse kidneys(*10*). We then determined cellular niches by clustering cells based on neighboring cell compositions. Uniquely in our study, we performed niche clustering separately within each renal compartment (cortex, OSOM and ISOM), thereby defining compartment-resolved microenvironments. In total, we identified 45 niches with distinct cellular compositions (14–18 niches per compartment; **Fig. 4, A to C**). Consistent with the previous observation(*10*), FR-PT-centered niches were associated with immune cells and fibroblasts in CKD. Remarkably, aTAL-centered niches, which were also enriched with immune cells and fibroblasts, were among the most abundant microenvironments in CKD across the cortex, OSOM, and ISOM. Moreover, aTAL-centered niches were consistently associated with M2-like macrophages across compartments (**Fig. 4, A to D**). Niches enriched with M2-like macrophages were associated with fibroblasts or myofibroblasts, supporting the pro-fibrotic signature of M2-like macrophages. To investigate intercellular signaling within aTAL niches, we performed ligand–receptor analysis using the snRNA-seq dataset. This analysis predicted multiple ligand–receptor pairs between aTAL cells and M2-like macrophages (**Fig. 4E**). Among these, IL-34, produced by tubular epithelial cells, has been implicated in renal fibrosis by promoting proliferation of intrarenal macrophages and bone marrow myeloid cells(*43*). Increased *Il34* expression in aTAL cells was verified by qPCR in primary TAL cells (**fig. S7F**). We also observed amyloid precursor protein (APP)-Cd74 interaction between aTAL cells and M2-like macrophages. This axis was recently reported as a key ligand-receptor pair between TAL cells and macrophages in an ADTKD-UMOD mouse model(*44*). These data suggest that aTAL cells form pathological niches with pro-fibrotic M2-like macrophages across compartments and could drive global kidney disease progression.

**Fig. 4.**
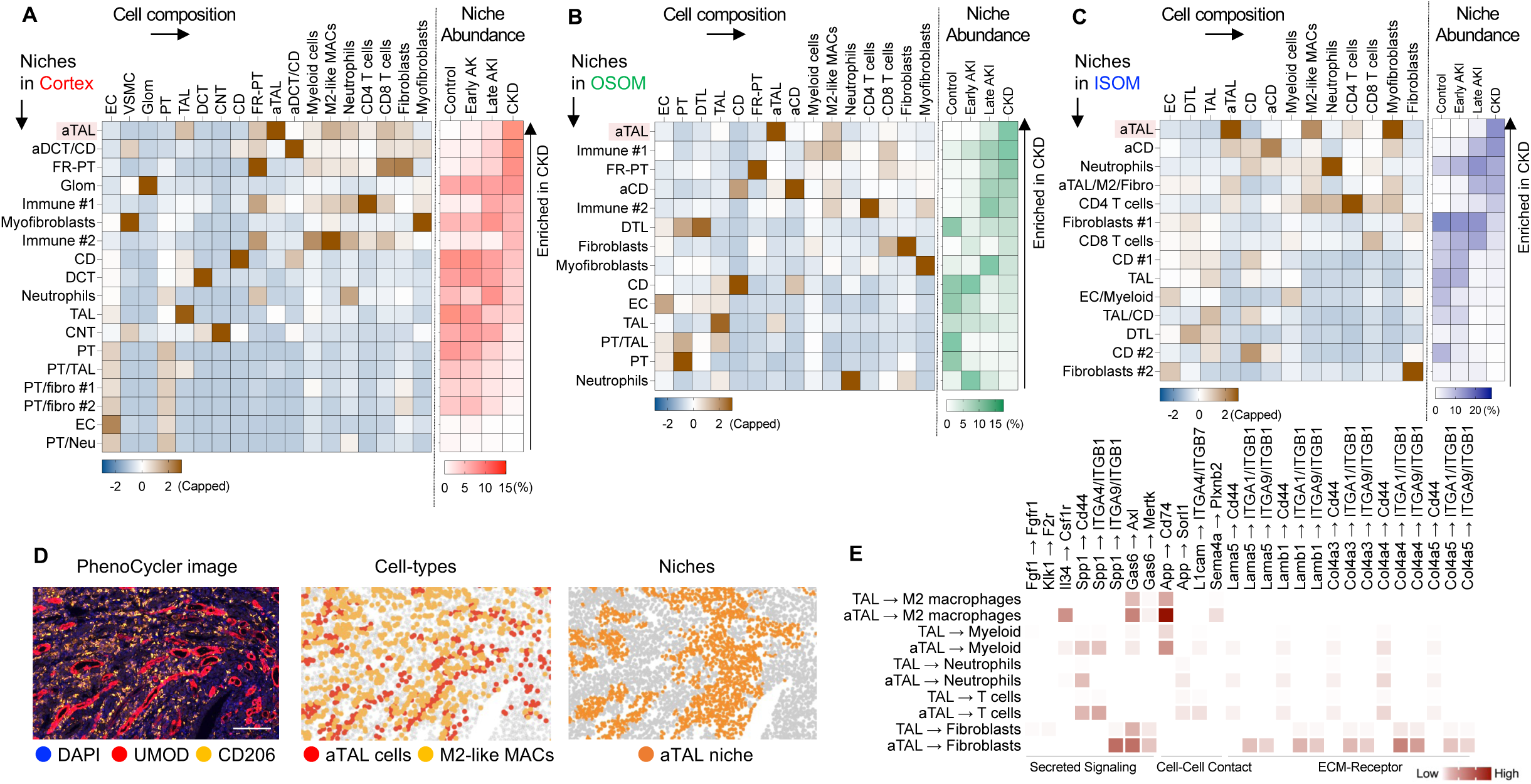
Altered TAL cells form CKD-specific niches with M2-like macrophages. (A-C) Compartment-resolved niche analysis in the IRI-induced CKD mouse model: (A) cortex, (B) OSOM, and (C) ISOM. Niches are ordered from top to bottom by their abundance in CKD. The left heatmap shows enrichment scores of cellular composition within each microenvironment. Enrichment scores were calculated by subtracting the regional frequency of each cell type from the cell composition within each niche, and were divided by the regional frequency of each cell type. Right heatmap showing the proportion of each niche in each timepoint. (D) Spatial distribution of marker proteins, cell types and niches in PhenoCycler dataset of IRI-induced CKD kidneys. Bar = 100 μm. (E) Putative ligand-receptor interactions predicted by CellChat in IRI-induced CKD kidneys.

### Reducing the burden of UMOD retention decreases aTAL cells and associated pro-fibrotic niches and mitigates the development of CKD

To determine whether intracellular UMOD retention sustains the pathological aTAL state and drives CKD development, we sought to suppress *Umod* expression during the window of AKI-to-CKD progression. Note that constitutional *Umod* knockout mice cannot be used in this experiment because they exhibit increased susceptibility to AKI(*16*), and differences in AKI severity are a major determinant of subsequent CKD progression(*45*). To this end, we generated a tamoxifen-inducible conditional *Umod* knockout (cKO) mouse. *Umod* knockdown starting day 1 after IRI (peak of AKI) suppressed *Umod* gene expression and prevented UMOD protein accumulation 4 weeks after IRI (**Fig. 5, A to C**). In this setting, wild-type (WT) mice developed CKD, as evidenced by impaired kidney function, fibrosis and global atrophy (**Fig. 5, D to F**). By contrast, *Umod* cKO mice showed markedly attenuated CKD development, with improved renal function and reduced fibrosis and atrophy (**Fig. 5, D to F**). To define the impact of UMOD retention on cell states and niche architectures, we performed PhenoCycler imaging on WT-IRI and cKO-IRI samples (n = 3 for each group; **Fig. 5G, fig. S8, A to C** and **table S2**). We identified 19 cell types, consistent with those observed in the IRI trajectory and IRI/AAN dataset. Note that UMOD expression, which is altered by the intervention, was not used to subtype TAL cells in this dataset. We found that aTAL cells were markedly reduced in cKO mice (**Fig. 5H**) and showed the greatest reduction among all cell types (**Fig. 5I**). To corroborate this finding, we next conducted snRNA-seq analysis (**fig. S9, A to C**). The snRNA-seq dataset showed 20 cell types, and aTAL cells were defined by Visium HD-derived label transfer (**fig. S9D and E**), using the same approach applied to the Control and CKD snRNA-seq dataset (**fig. S3D and E**). In the snRNA-seq dataset, aTAL cells were reduced in cKO mice to a similar extent as in the PhenoCycler dataset (**fig. S9F**). These results indicate that altered TAL states are driven by the burden of UMOD retention. The concomitant reduction in the abundance of M2-like macrophages in the *Umod* cKO PhenoCycler data (**Fig. 5I**) suggests a functional link between aTAL abundance and M2-like macrophage accumulation.

**Fig. 5.**
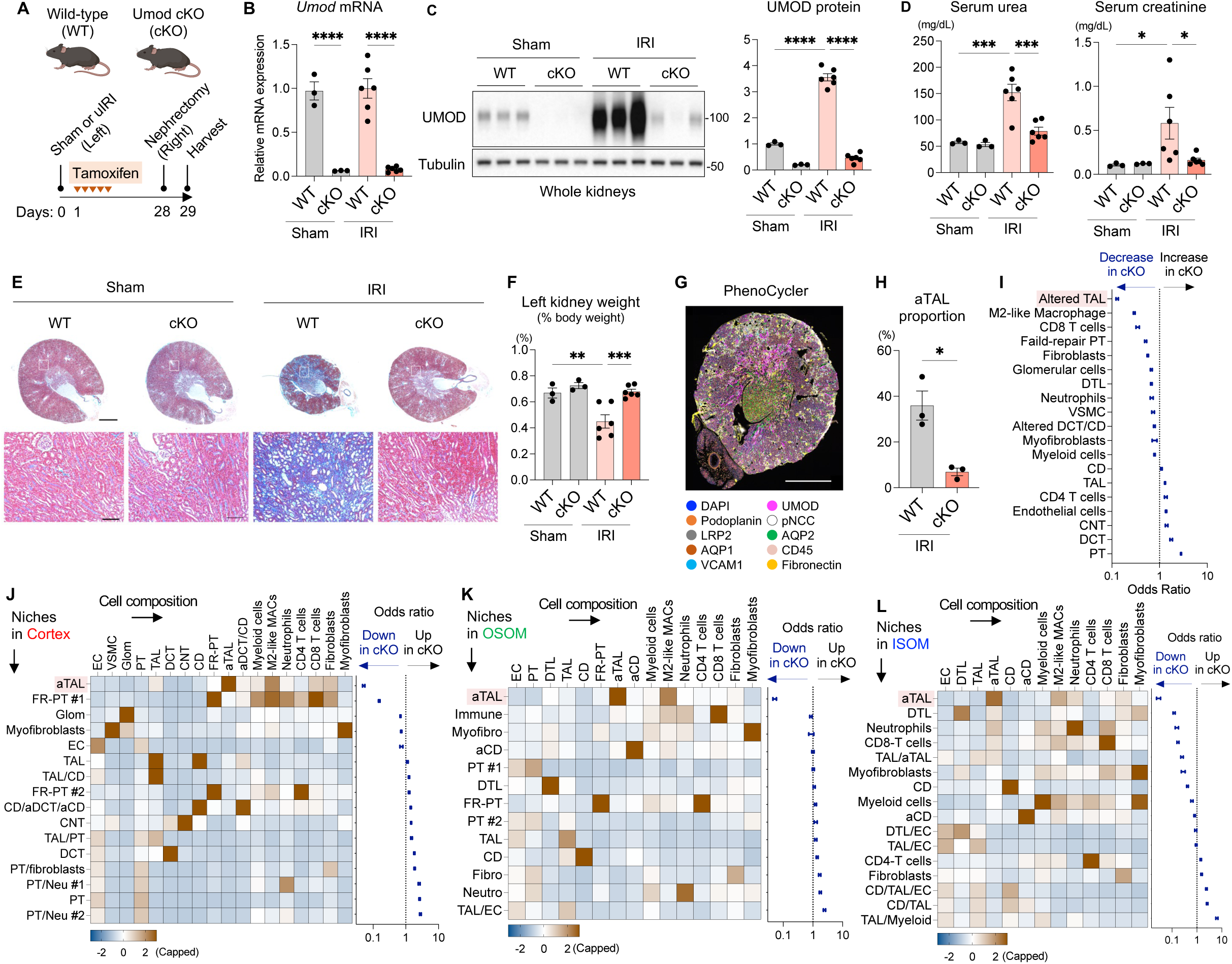
Reducing the burden of UMOD retention decreases aTAL cells and associated pro-fibrotic niches and mitigates the development of CKD. Wild-type (WT) and conditional Umod knockout (cKO) mice underwent sham or unilateral IRI. These mice received tamoxifen for consecutive 5 days starting on IRI day 1. Mice were harvested 4 weeks after IRI. Contralateral nephrectomy was performed 24 hours before harvest to assess kidney function. (A) Schematic of the experimental design. (B) Relative *Umod* mRNA expression normalized to *Gapdh*. (C) Immunoblotting of kidney UMOD. (D) Serum urea and creatinine levels. (E) Masson’s trichrome staining of IRI kidneys. Bars = 1 mm or 100μm (zoom in). (F) Left kidney weight as percent of baseline body weight. n = 3 for Sham WT and Sham cKO; n = 6 for IRI WT and IRI cKO for panels b-f. (G-L) PhenoCycler protein imaging was conducted on WT-IRI and cKO-IRI samples (n = 3 per genotype). (G) Representative PhenoCycler multiplexed protein image. Bar = 1 mm. (H) Proportion of aTAL cells among TAL cells in each genotype. (I) Odds ratios of cell-type abundance by genotype. (J-L) Niche analysis of WT-IRI and cKO-IRI kidneys: (J) cortex, (K) OSOM, and (L) ISOM. Niches are ordered from top to bottom by their reduction in cKO relative to WT. The left heatmap shows enrichment score of cellular composition within each microenvironment. Enrichment scores were calculated by subtracting the regional frequency of each cell type from the cell composition within each niche, and were divided by the regional frequency of each cell type. Right panel showing the odds ratios of niche abundance by genotype. Values are represented as mean ± SEM. *p<0.05, **p<0.01, ***p < 0.001, ****p < 0.0001.

To link these cell-level alterations to whole-kidney disease progression, we performed niche analysis (**Fig. 5, J to L**) using the approach used for the IRI trajectory dataset. Genotype-based comparison of niche abundance revealed that the aTAL niche, consistently enriched for M2-like macrophages, showed the greatest reduction in cKO across all regions. These data suggest that aTAL-M2-like macrophage microenvironments are key drivers of CKD progression. In contrast, FR-PT niches were minimally affected by cKO; the cortical FR-PT #1 niche showed a modest reduction, whereas the cortical FR-PT #2 and OSOM FR-PT niches were not decreased. These data further support that *Umod* cKO specifically reduces aTAL niches rather than disrupting global epithelial-immune crosstalk. Taken together, the data support that UMOD retention drives pathogenic TAL cell states and associated profibrotic niches, which lead to CKD progression.

### Conserved altered TAL cell state in human kidneys

To determine the clinical relevance of our findings, we analyzed a publicly available human kidney PhenoCycler dataset from the Kidney Precision Medicine Project consortium (**Fig. 6A**) that was used in a recent publication(*46*). We analyzed 1,138,996 cells from 36 individuals (healthy reference tissue, n = 8; AKI, n = 12; CKD, n = 16; patient demographics are shown in **table S3**). Note that renal biopsies from AKI patients are typically obtained beyond the earliest phase of injury and therefore likely to correspond to late AKI rather than early AKI stage when contextualized within the experimental AKI-to-CKD transition. Cell clustering identified 18 distinct clusters (**Fig. 6, B and C** and **fig. S10A**). In this dataset, neutrophils are presumed to be included within the “Myeloid cells” cluster due to the lack of specific markers. CD206-high macrophages may include pro-fibrotic M2-like macrophages described in mice. However, in human kidneys, CD206 is also an established marker of resident homeostatic macrophages(*47*). Therefore, we annotated these cells as CD206-high macrophages, likely representing a mixture of homeostatic and pro-fibrotic populations.

**Fig. 6.**
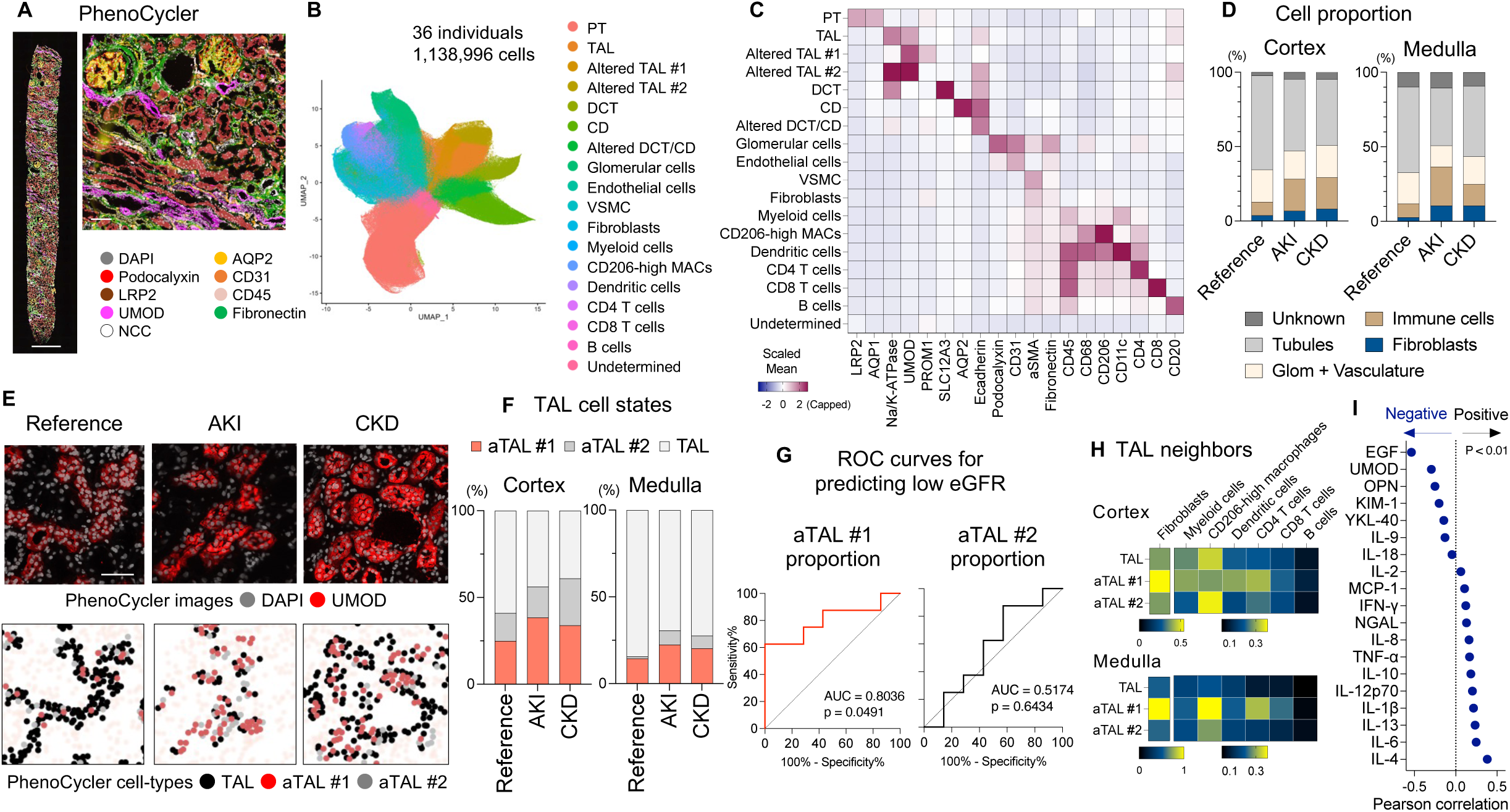
Conserved altered TAL cell state in human kidneys. Human kidney PhenoCycler datasets from the Kidney Precision Medicine Project consortium were analyzed. (A) Representative PhenoCycler multiplexed protein image. Bars = 1mm or 100 μm (zoom in). (B) UMAP plot showing cell types. (C) Heatmap displaying mean expression of marker proteins across clusters. (D) Cellular composition at each disease state. (E) Spatial distribution of TAL subtypes and corresponding UMOD staining of PhenoCycler dataset on human kidneys. Bars = 50 μm. (F) Proportion of altered TAL within total TAL cells in each timepoint. (G) ROC analysis of aTAL cell proportions for predicting low eGFR patients. The proportions of altered TAL #1 and altered TAL #2 within TAL cells in the cortex were calculated for each CKD patient (n = 15). Low eGFR was defined as the lower 50% of eGFR values among CKD patients, and the ability of altered TAL proportions to predict low eGFR was assessed. (H) Neighborhood analysis of TAL subtypes. For each TAL cell, the number of neighboring fibroblasts and immune cells within 30 μm was counted. (I) Pearson correlation coefficients between cortical aTAL #1 cell proportion and urinary biomarkers measured by multiplex immunoassays. Among 34 individuals with available cortical PhenoCycler data, 26 individuals with multiplex immunoassay data were included in the analysis.

Cell composition analysis showed that AKI and CKD were marked by increased immune cell and fibroblast populations in both cortex and medulla (**Fig. 6D**). When examined in detail, fibroblasts and immune cell subtypes increased in disease with distinct spatial and temporal distribution (**fig. S10B**). In the cortex, CD206-high macrophages were already abundant in reference tissues, suggesting that the resident homeostatic population is predominant. In contrast, in the medulla, CD206-high macrophages were disease-specific, consistent with M2-like macrophages described in mice.

We detected two altered TAL populations (aTAL #1 and aTAL #2) with higher UMOD expression compared to healthy TAL cells (**Fig. 6C**). aTAL #1 exhibited features of dedifferentiation (decreased Na/K-ATPase and E-cadherin expression and increased PROM1 expression), closely resembling the aTAL population identified in the mouse IRI model. In contrast, aTAL #2 showed the highest Na/K-ATPase expression. This subtype was not identified in the mouse IRI model and likely represents human-specific transitional, adaptive or alternative cell states. These two altered TAL cell populations were increased in disease (**Fig. 6, E and F**). These altered populations were also present in the reference specimens, consistent with prior observations of altered tubules in healthy human kidneys(*13*, *48*). Notably, the proportion of aTAL #1, but not aTAL #2, in CKD specimens predicted reduced kidney function at the time of biopsy (**Fig. 6G**). Further, aTAL #1 was specifically associated with neighboring fibroblasts and immune cells (**Fig. 6H**), suggesting a role in fibro-immune microenvironments. The exception was cortical CD206-high macrophages, which are likely homeostatic populations as noted above. Collectively, our findings show that in humans, aTAL #1 population, which closely resembles mouse aTAL cells, is associated with impaired kidney function and fibro-immune microenvironments, suggesting a conserved pathological role of TAL states in kidney diseases.

Urinary UMOD levels are known to be decreased in patients with CKD(*16*, *49*), consistent with intracellular UMOD retention. We examined the correlation between aTAL #1 proportion and urinary biomarker levels for 26 individuals for whom both PhenoCycler imaging data and urinary multiplexed immunoassay data were available (**Fig. 6I**). Among 19 biomarkers measured by the multiplex assay, urinary EGF concentration showed the strongest negative correlation with the proportion of aTAL #1 cells. Given that EGF was among the most downregulated genes in murine aTAL cells (**fig. S3G**) and that intra-kidney EGF gene expression is highly correlated with urinary EGF levels(*34*), decreased urinary EGF may serve as a potential surrogate marker of aTAL states. Unsurprisingly, urinary UMOD levels showed the second-strongest negative correlation (albeit not statistically significant, likely due to sample size), consistent with impaired urinary UMOD secretion in altered TAL state in humans.

## DISCUSSION

Emerging studies have identified injury-induced kidney tubular epithelial cell states, particularly for the PT epithelium, which persist beyond acute injury and into CKD. How these altered cell populations drive CKD development remains undetermined. Here, we identified a disease-specific inflammatory aTAL cell state that forms profibrotic microenvironments driving CKD progression. Mechanistically, aTAL state was driven by intracellular retention of UMOD, as genetic reduction of UMOD burden diminished aTAL abundance and associated profibrotic microenvironments, and prevented CKD development. These findings suggest a key role for altered TAL states in the pathogenesis and progression of CKD. The conservation of this phenotype in human kidney disease and its association with reduced kidney function underscore its potential clinical relevance. In particular, targeting maladaptive tubular cell states may represent a promising therapeutic strategy because these states lie upstream of inflammation and fibrosis and could provide an opportunity to address a complementary mechanism and further reduce progressive kidney damage beyond current therapies for CKD.

This study highlights that spatial proteomics can define key features of disease-associated cell states that are not resolved by transcriptomic profiling. Although recent single-cell and spatial transcriptomic studies have substantially advanced our understanding of injury-associated epithelial cell states, transcript abundance does not necessarily reflect protein localization and intracellular processing, key determinants of cellular function and disease mechanisms. Here, spatial proteomics revealed intracellular UMOD accumulation as a key feature of altered TAL states. More broadly, our findings demonstrate that protein-level alterations can redefine disease-associated cell states beyond transcript abundance, providing an additional layer for understanding disease biology.

By demonstrating that intracellular UMOD retention drives disease through perpetuation of TAL dysfunction, our results help resolve a longstanding paradox in UMOD biology. Genetic variants associated with increased UMOD expression confer a higher risk of CKD. However, UMOD itself has important protective functions when appropriately secreted, including preventing urinary tract infections and nephrolithiasis through its apical release into the urine, and modulating immune responses and systemic oxidative stress through its minor basolateral release into the kidney interstitium and circulation(*16*). Consistent with these protective effects, urinary UMOD levels decline in kidney disease, and higher urinary UMOD levels are associated with favorable renal outcomes(*16*, *49*). Our findings provide a mechanistic framework that reconciles these observations. Following kidney injury, increased UMOD expression in TAL cells with impaired secretory machinery may increase intracellular UMOD retention, promoting a persistent dysfunctional TAL state and accelerating CKD progression. Consistent with this model, a recent clinical study showed that lower urinary UMOD levels following AKI are associated with subsequent CKD progression(*50*). In this context, impaired UMOD processing and secretion uncouple UMOD production from its release, explaining why genetic variants that increase UMOD expression can increase CKD risk despite the apparent protective association of higher urinary UMOD levels. Accordingly, our data suggest that low urinary UMOD levels largely reflect impaired secretion and increased burden of dysfunctional aTAL cells with intracellular UMOD retention, rather than simply reduced UMOD production.

Our data also suggest that urinary EGF levels may reflect TAL states, which is consistent with reports indicating that urinary EGF is an emerging CKD biomarker(*33*, *34*). Future studies are needed to determine whether changes in urinary UMOD and EGF levels could reflect underlying TAL states, potentially enabling biomarker-guided patient stratification and precision medicine approaches for CKD therapies.

Our findings also suggest that intracellular UMOD retention may represent a shared pathological feature across common and rare forms of CKD. UMOD has long been implicated in kidney disease, with common non-coding variants associated with CKD risk and rare coding mutations causing monogenic CKD through protein misfolding. Here, we show that intracellular UMOD retention extends beyond monogenic disease to common CKD, where it arises from impaired secretory machinery rather than protein misfolding. These results suggest that therapeutic strategies aimed at reducing intracellular UMOD retention may be broadly applicable across diverse forms of CKD. Whether restoring UMOD processing and secretion is sufficient to reverse pathological TAL states and CKD progression remains an important question for future investigation.

In conclusion, we identify a conserved inflammatory aTAL state defined by intracellular UMOD retention that promotes CKD progression. These findings may provide a mechanistic explanation for longstanding observations in UMOD genetics and biomarker studies, while highlighting UMOD retention and maladaptive tubular cell states as potential therapeutic targets in CKD.

## MATERIALS AND METHODS

### Animal experiments

We used 8-to 12-week-old C57BL/6J wild-type mice (The Jackson Laboratory) and newly generated conditional *Umod* knockout mice (described below). All animals were maintained in a temperature-controlled room with free access to food and water and a 12-hour day/night cycle. IRI was performed by left unilateral renal pedicle clamping. Sham surgery was done by the same procedure without clamping. Animals were anesthetized with isoflurane. Induction occurred with 3% vapor at 1.5 l/min flow of oxygen. Animals were maintained on a rodent anesthesia circuit at 1-2% vapor at 1.5 l/min flow of oxygen. Core temperature was maintained from 36.5-37.0°C utilizing a homeothermic warming pad. Daily saline supplementation was conducted after surgery. For aristolochic acid nephropathy model, a single dose of aristolochic acid Ⅰ (A5512, Sigma-Aldrich) dissolved in DMSO, was injected via intraperitoneal injection. For 5/6 nephrectomy model, two-thirds of the left kidney was resected, followed by right nephrectomy 1 week later. Serum creatinine concentration was measured by liquid chromatography-tandem mass spectrometry by the University of Alabama O’Brien Center. Serum urea concentration was measured using QuantiChrom Urea Assay Kit (DIUR100, BioAssay Systems).

### Histology

Mouse kidney tissues were collected and fixed in 4% PFA and embedded in paraffin. Paraffin embedding, tissue sectioning and Masson’s Trichrome staining were performed by the Histology Lab Service Core in Indiana University. Images were taken by Keyence BZ-810.

### PhenoCycler multiplexed imaging of mouse kidney FFPE tissues

PhenoCycler (formerly CODEX; Co-Detection by Indexing) is an imaging-based spatial protein profiling methodology(*51*, *52*). The latest PhenoCycler-Fusion (PCF) 2.0 platform (Akoya Biosciences) was used as described recently(*46*, *49*), with modifications to accommodate FFPE tissues instead of OCT-embedded samples. Mouse FFPE kidney sections (5 μm thickness) were cut using a microtome and mounted on Superfrost Plus Gold slides (Thermo Fisher Scientific, #1518848). Slides were deparaffinized in xylenes and graded ethanol, followed by antigen retrieval in citrate buffer using a pressure cooker (115 °C, 14 min) according to the Akoya PhenoCycler protocol. After washing and equilibration in hydration and staining buffers, tissues were incubated overnight at 4 °C with a PhenoCycler antibody cocktail. Iterative oligonucleotide probe staining and automated tile-scan imaging were performed using the PCF fluidics handler and integrated microscope. Images were processed as part of the PCF imaging workflow.

### PhenoCycler image analysis of mouse kidney FFPE tissues

PhenoCycler image analysis was conducted as described previously(*46*), with modifications. Final qptiff images were imported into QuPath v0.6.0 to crop biopsies into individual image files. Nuclear segmentation was conducted using ImageJ plugin VTEA based on 4′,6-diamidino-2-phenylindole (DAPI) staining, and cells were defined using the Ring6C perinuclear expansion generated from nuclear segmentation. Mean intensities of each protein within cells were quantified. One cortical and three medullary compartments were defined for each tissue based on the expression of region-specific markers. Perirenal adipocytes were not included in the analysis. These cell-by-marker intensity matrices were combined and imported into R. Cells with the top 1% of DAPI signal were excluded for each sample. After filtering, the mean intensities of proteins were z-scaled. VWF, CD11c, Ly6c, Vimentin, PLSCR1 and Collagen1A were excluded from analyses due to insufficient signal-to-noise ratio or specificity. Parental clustering was performed using the Louvain-based FastPG method and a feature space including scaled expression of LRP2, AQP1, UMOD, pNCC, Calbindin, AQP2, E-cadherin, Na/K-ATPase, aSMA, Fibronectin, CD45, CD74 and CD206. Uniform manifold approximation and projection (UMAP) embeddings were also calculated. Clusters were annotated based on marker expression and spatial localization by mapping the clusters onto specimens. For IRI trajectory dataset, PT/DTL/DVR, TAL, and immune cell clusters were reclustered separately using key marker features (PT/DTL/DVR: LRP2, AQP1, VCAM1, CD31; TAL: UMOD, Na/K-ATPase; immune cells: CD74, CD206, Ly6G, CD153, CD3, CD8a, fibronectin). For IRI and AAN dataset, PT, immune, and VSMC/myofibroblasts clusters were reclustered separately using key marker features (PT: LRP2, AQP1, VCAM1; immune cells: CD74, CD20, Ly6G, CD153, CD3, CD4, CD8a, B220, fibronectin; VSMC/myofibroblasts: fibronectin, aSMA, CD31). For conditional *Umod* knockout mouse dataset, TAL, immune cell, and VSMC/myofibroblasts clusters were reclustered separately using key marker features (TAL: Na/K-ATPase; immune cells: CD74, CD206, Ly6G, CD153, CD3, CD8a, fibronectin; VSMC/myofibroblasts: fibronectin). For reclustering, FastPG was applied when using two or more features, whereas k-means clustering was used for single-feature analyses. Clusters with highly similar expression profiles were carefully merged. DVR populations were included in endothelial cell clusters for the downstream analysis.

### Cellular neighborhood analysis of mouse kidneys

Identification and quantification of mouse kidney niches were performed using the PhenoCycler dataset separated by regions. For each cell, neighboring cells within a 30 μm centroid-to-centroid distance were counted. These counts were converted to proportions and mean-centered within each cell type. The neighborhood features were extracted and subjected to principal component analysis (PCA), followed by FastPG clustering to define neighborhood clusters. Clusters were annotated based on the enriched cell types. Clusters with highly similar cell type compositions were carefully merged.

### PhenoCycler image analysis of human kidney tissues

This study used publicly available, de-identified human PhenoCycler datasets from our previous study from the Kidney Precision Medicine Project (KPMP) consortium(*46*). Specimens were acquired with informed consent and approved under a protocol by the KPMP single Institutional Review Board (IRB) of Washington University in St. Louis (Approval No. 20190213). In this study, biopsy specimens with adjudicated diagnoses(*49*) were selected, excluding those with “cannot be determined”. Cortex (n = 34) and medulla (n = 11) specimens derived from 36 patients (healthy reference tissues (HRT), n = 8; AKI, n = 12; CKD, n = 16) were used for subsequent analyses. Image analysis was conducted using a pipeline similar to that applied to the mouse dataset. Parental clustering was performed based on the expression of CD45, NCC, AQP1, fibronectin, CD31, Na/K-ATPase, E-cadherin, Podocalyxin, LRP2, αSMA, AQP2, UMOD, CD68, and CD206. Immune cell clusters were subsequently reclustered using the expression of CD8, CD3, CD45RO, CD20, CD4, CD45, CD11c, FOXP3, CD68, CD206, and HLA-DR. Cluster annotation was performed based on marker expression and spatial localization by mapping the clusters onto specimens.

### Visium HD assay

Visium HD Spatial Gene Expression (Visium HD) is a whole-transcriptome assay with single-cell-scale resolution with a markedly increased oligonucleotide barcode density (∼11,000,000 continuous 2-μm features in a 6.5 × 6.5-mm capture area)(*53*). To ensure RNA quality, we first isolated bulk RNA from FFPE sections using RNeasy FFPE Kit (73504, Qiagen) and assessed DV200 by RNA ScreenTape. Serial sections (5 μm thickness) for PhenoCycler analysis were deparaffinized, stained with H&E and decrosslinked according to the Visium HD Spatial Gene Expression for FFPE protocol (CG000684, 10x Genomics). H&E-stained tissues were imaged on a Keyence BZ-810 using a ×20 objective. Immediately after decrosslinking, libraries were prepared according to the user guide for Visium HD Spatial Gene Expression Reagent Kits (CG000685, 10x Genomics). Final libraries were sequenced using Illumina NovaSeq X Plus platform using the 10B flow cell with paired-end reads.

### Visium HD analysis

Raw sequencing data were processed using Space Ranger v4. Cell segmentation was performed by identifying nuclei from high-resolution H&E images, followed by Voronoi-like tessellation to approximate cell boundaries, as implemented in Space Ranger. Downstream data analysis was performed using R and Seurat v5 package. Seurat objects were generated and cell-level polygon area and centroid spatial coordinates were incorporated. Cells were filtered by excluding the top and bottom 1% of polygon areas within each sample. Cells were further filtered to retain those with ≥50 detected genes and ≥100 total transcripts per cell. Each sample was normalized using SCTransform. Datasets were then integrated using SCT-based anchor identification, followed by dimensionality reduction (PCA and UMAP) and graph-based clustering. Clusters were annotated based on gene expression profiles and spatial localization by mapping them to specimens.

### PhenoCycler - Visium HD spatial registration

Landmark-based registration was performed to co-register PhenoCycler and Visium HD images. Corresponding anatomical landmarks in the PhenoCycler images and the H&E images associated with the Visium HD dataset were selected using ImageJ plugin BigWarp. Coordinates of PhenoCycler cells were then transformed into the Visium HD coordinate space using thin-plate spline (TPS) interpolation using R. For each PhenoCycler cell, the nearest Visium HD cell was assigned. The distance to the nearest cell was within 5 μm for 62.9% of cells and within 10 μm for 92.9% of cells. Cells with a nearest-neighbor distance ≤ 10 μm were used for the subsequent analysis. Cell-type annotations were compared between the two modalities and enrichment scores were calculated to adjust for differences in cell-type frequencies. In detail, the proportion of Visium HD annotations assigned to each PhenoCycler cell was subtracted by the overall frequency of each cell type in the Visium HD sample, and then divided by this overall frequency.

### GEM-X Flex Gene Expression assay

GEM-X Flex Gene Expression assay measures RNA levels at single cell/nucleus levels in fixed samples using probes targeting the whole coding transcriptome. RNA quality was first assessed using the DV200 metric, as in the Visium HD workflow. For all the experiments, kidneys from three animals per condition were pooled. Nuclei from FFPE tissues were isolated according to the Sample Preparation from FFPE Tissue Sections for GEM-X Flex Gene Expression protocol (CG000784, 10x Genomics). Following probe hybridization and ligation, samples were loaded onto the Chromium Controller (10x Genomics) to generate Gel Beads-in-Emulsion (GEMs) using the GEM-X technology with a target recovery of 20,000 cells per sample. After GEM incubation, emulsions were broken and barcoded products were recovered, followed by PCR amplification and library construction according to the manufacturer’s protocol. Sequencing libraries were generated and sequenced on an Illumina Novaseq X plus system using the 10B flow cell with paired-end reads.

### GEM-X Flex Gene Expression analysis

Raw sequencing data were processed using Cell Ranger v9. Data analysis was conducted using R and Seurat v5 package. Seurat objects were generated and cells were filtered to retain those with ≥ 300 and ≤ 7,999 detected genes, and ≤ 10% mitochondrial transcripts. Ambient RNA was then corrected using SoupX and doublets were subsequently identified and removed using scDblFinder. Each sample was normalized using SCTransform. Datasets were then integrated using SCT-based anchor identification, followed by dimensionality reduction (PCA and UMAP) and graph-based clustering. Clusters were annotated based on gene expression profiles. Visium-derived TAL labels (healthy TAL (TAL #1 and TAL #2) and aTAL) were transferred to snRNA-seq TAL cell clusters using anchor-based integration (FindTransferAnchors and TransferData functions). Subsequent analyses were conducted using TAL cells with prediction scores ≥ 0.7. Differentially expressed genes were determined using FindMarkers function with the Wilcoxon rank-sum test. Gene set enrichment analysis was performed using fgsea with Hallmark pathways. Ligand-receptor interaction analysis was conducted using CellChat package.

### Immunoblotting

Mouse kidneys were homogenized in Pierce IP Lysis Buffer (87787, Thermo Fisher Scientific) with Halt Protease and Phosphatase Inhibitor Single-Use Cocktail (78442, Thermo Fisher Scientific). The lysates were centrifuged at 600xg for 10 minutes at 4°C to remove debris, and the supernatants were collected. Tissue lysates and urine were denatured for 20 minutes at 60°C using Sample Buffer, Laemmli 2× Concentrate (S3401, Sigma). Total protein concentration was determined using Detergent Compatible Protein Assay (5000113-5000115, BioRad). For immunoblotting, the following primary antibodies were used: rabbit anti-UMOD (1:5000, ab207170, Abcam), rabbit anti-spliced XBP1 (1:2000, 40435, Cell Signaling Technology), rabbit anti-ATF4 (1:2000, 11815, Cell Signaling Technology) and rabbit anti-α-tubulin (1:2000, 11224-1-AP, Proteintech). HRP-conjugated anti-IgG antibodies (Proteintech) were used as secondary antibodies. Coomassie brilliant blue staining was conducted using One-Step Blue® Protein Gel Stain (21003-1L, Biotium). Loading controls were run on the same membrane as the target protein after stripping using Restore Western Blot Stripping Buffer (21059, Thermo Fisher Scientific). The signals were quantified by ImageJ.

### Quantitative RT-PCR (qPCR)

Mouse kidneys were homogenized in TRIzol Reagent (15596018, Thermo Fisher Scientific). Total kidney RNA was isolated following the manufacturer’s instructions. cDNA was synthesized using ReverTra Ace (TYB-FSQ-201, Diagnocine). qPCR analysis was performed on the Thermal Cycler Dice Real Time System using TaqMan Gene Expression Master Mix (4369016, Thermo Fisher Scientific) or THUNDERBIRD Next SYBR qPCR Mix (TYB-QPX-201, Diagnocine). *Gapdh* mRNA levels were used for normalization. For TaqMan qPCR, we used the following TaqMan probes: mouse *Umod* (Mm01137672_m1, Thermo Fisher Scientific) and mouse *Gapdh* (4352339E, Thermo Fisher Scientific). For SYBR qPCR, we used custom mouse *AS-Umod* and *Gapdh* primers as described previously(*39*).

### Immunostaining

Mouse kidneys were fixed with 4% paraformaldehyde and cut into 50 μm sections using vibratome (Leica). Mouse kidney sections were incubated with the following primary antibodies: sheep anti-UMOD (AF5175, R&D Systems), rabbit anti-NKCC2 (SPC-401, StressMarq Biosciences), rabbit anti-CANX (C4731, Sigma) and rabbit anti-GM130 (70767, Cell Signaling Technology). Alexa Fluor 488 and 647 were used as the secondary antibodies. The stained sections were mounted with ProLong Glass (P36982, Thermo Fisher Scientific). Immunofluorescence images were obtained using SP8 Confocal Microscope (Leica).

### Isolation of primary TAL cells

Primary TAL cells were isolated by magnetic cell separation as described previously(*39*, *54*). Mouse kidneys were decapsulated, minced, and then digested in a digestion buffer (2 mg/ml Collagenase type I (SCR103, Sigma-Aldrich) and 100 U/ml DNase (79254, Qiagen) in serum-free DMEM/F12 (11320033, Thermo Fisher Scientific)) at 37 °C shaker for 60 min. Then, DMEM/F12 with 10% FBS was added to halt the enzymatic digestion. Single-cell suspension was collected using a 40 μm cell strainer (40ICS, MIDSCI). Next, Dynabeads Biotin Binder (11047, Thermo Fisher Scientific) was conjugated with mouse anti-UMOD Biotinylated Antibody (BAF5175, R&D Systems). After washing, the beads were incubated with the single-cell suspension at 4 °C for 60 min. Both UMOD-positive cells (on-beads) and UMOD-negative cells (supernatant) were washed and collected as TAL cells and non-TAL cells, respectively.

### Generation of conditional Umod knockout mice

An 11.25 kb genomic region encompassing Umod exons 5–6 was subcloned from a C57BL/6 BAC and engineered to introduce a LoxP site 253 bp upstream of exon 5 and a LoxP/FRT-flanked neomycin cassette 300 bp downstream of exon 6. The targeting construct, containing homology arms, was linearized and electroporated into C57BL/6 FLP embryonic stem cells. Correctly targeted clones were injected into BALB/c blastocysts to generate chimeric mice. Germline transmission was achieved by crossing to C57BL/6 mice. Umod*^flox/flox^* mice were crossed with Rosa26-CreERT2 mice (The Jackson Laboratory, #008463) to generate tamoxifen-inducible *Umod* knockout mice. *Umod*^flox/flox^;Rosa26-CreERT2 mice were intercrossed, and littermate mice lacking Cre were used as controls (defined as wild-type in this manuscript), and mice homozygous for Cre were used as conditional *Umod* knockout (*Umod* cKO) mice. To induce gene deletion, 75 mg/kg of tamoxifen (Sigma, #T5648) dissolved in corn oil was administered intraperitoneally for 5 consecutive days starting 1 day post-surgery. Based on preliminary experiments showing that maximal reduction of UMOD protein levels was achieved 2 weeks after tamoxifen administration, mice were harvested 4 weeks after IRI to allow robust evaluation of phenotypic differences resulting from UMOD depletion.

### Urinary biomarker measurement

We assessed publicly available urinary biomarker immunoassay data, generated by the Kidney Precision Medicine Project (Accessed Sep 18th, 2024, https://www.kpmp.org). Urine handling and processing were conducted following the KPMP Biospecimen Manual of Procedures (MOP). Urinary IFN-γ, IL-1β, IL-2, IL-4, IL-6, IL-8, IL-10, IL-12p70, IL-13, and TNF-α concentrations were measured using the Meso Scale Discovery (MSD) V-PLEX Proinflammatory Panel 1 kit; IL-9 concentrations were measured using the MSD V-PLEX Cytokine Panel 2 kit; and NGAL, UMOD, OPN, EGF, YKL-40, IL-18, MCP-1, and KIM-1 concentrations were measured using a custom MSD multiplex assay. All data and quality control results are available at the Data Repository of the KPMP website (https://atlas.kpmp.org/repository/?size=n_20_n).

### Sex as a biological variable

This study exclusively examined male mice. Since UMOD expression levels and susceptibility to IRI differ between sexes(*55*, *56*), the extent to which our findings are applicable to females remains undetermined. For patient samples, we included both males and females, but the sample size was not enough to assess sex-specific differences. We recognize the importance of considering sex as a biological variable and plan to address this question in future studies.

### Statistics and reproducibility

Statistical analyses were performed in Excel and GraphPad Prism and quantitative data were presented as means ± SEM unless otherwise indicated. Statistical significance was evaluated using an unpaired t-test (between two conditions) or one-way ANOVA with embedded comparisons (among multiple conditions). A two-sided p < 0.05 was considered statistically significant. For animal experiments using conditional knockout mice, investigators were blinded to genotype.

## Acknowledgements

Histology work was supported by the Histology Core of the Indiana Center for Musculoskeletal Health at IU School of Medicine and the Indiana Clinical Translational Sciences Institute (CTSI). Sequencing analysis was carried out in the Center for Medical Genomics at Indiana University School of Medicine, which is partially supported by the Indiana University Grand Challenges Precision Health Initiative. We gratefully acknowledge the essential contributions of the patient participants and the support of the American public through their tax dollars. The content is solely the responsibility of the authors and does not necessarily represent the official views of the National Institutes of Health.

## Funding

National Institute of Health, National Institute of Diabetes and Digestive and Kidney Diseases R01DK146350 (TME)

VA Merit Award 5I01BX003935 (TME)

Dialysis Clinic Inc (TME)

Simon Foundation grant (TME)

American Society of Nephrology Kidney Cure (TME)

Takeda Science Foundation (AN)

American Heart Association Postdoctoral Fellowship https://doi.org/10.58275/AHA.26POST1556544.pc.gr.240432 (AN)

Ralph W. and Grace M. Showalter Research Trust and the Indiana University School of Medicine (DB)

National Institute of Health, National Institute of Diabetes and Digestive and Kidney Diseases

U01DK133081, U01DK133091, U01DK133092, U01DK133093, U01DK133095, U01DK133097, U01DK114866, U01DK114908, U01DK133090, U01DK133113, U01DK133766, U01DK133768, U01DK114907, U01DK114920, U01DK114923, U01DK114933, U24DK114886, UH3DK114926, UH3DK114861, UH3DK114915, and UH3DK114937 (KPMP).

## Author contributions

Conceptualization: AN

Methodology: AN, ARS, TME

Investigation: AN, GJR, WSB, SK, MA, DLW, DB, TK, AT, HS, AK, KAL

Visualization: AN, TME

Funding acquisition: AN. DB, TME

Project administration: AN, TME Supervision: TME

Writing – original draft: AN

Writing – review & editing: ACR, JPL, KRT, PCD, JH, LR, TME

## Competing interests

The authors declare they have no competing interests.

## Data availability

Raw and processed data of PhenoCycler, Visium HD and snRNA-seq experiments will be available in a public repository upon acceptance of the peer-reviewed manuscript. De-identified participant-level data, including PhenoCycler multiplex imaging data and urinary immunoassay data, are publicly available in the KPMP website (www.KPMP.org).

## Code Availability

Code to reproduce figures will be available upon acceptance of the peer-reviewed manuscript. No additional custom computational code was generated in this study.

## Supplementary Materials

Figs. S1 to S10

Tables S1 to S3

Data S1 and S2

## Notes

### Competing Interest Statement

The authors have declared no competing interest.

