## Supplementary Materials for "Spatial proteomics identifies altered thick ascending limb cells as drivers of kidney disease through uromodulin retention"

Azuma Nanamatsu *et al.*

Corresponding authors:
Azuma Nanamatsu, and Tarek M. El-Achkar,

**The PDF file includes:**

Figs. S1 to S10
Tables S1 to S3

**Other Supplementary Materials for this manuscript include the following:**

Data S1 to S2

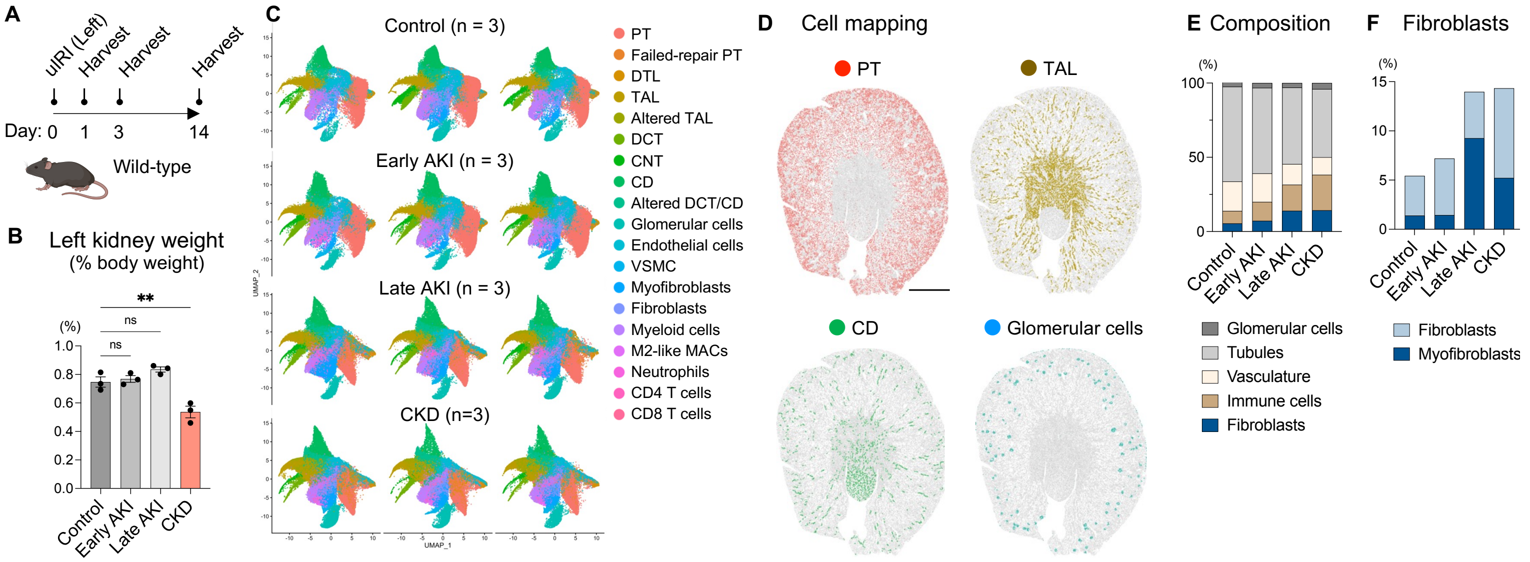


**Fig. S1** (relating to Fig. 1): (A) Schematic of the experimental design. Wild-type mice underwent unilateral IRI with 23-minute clamp time and were harvested at multiple time points. (B) Left kidney weight as percent of baseline body weight at each disease stage. (C) UMAP plots of all individual samples. (D) Cellular distribution in control kidneys. Bar = 1 mm. (E) Cellular composition at each disease stage. (F) Proportion of fibroblasts and myofibroblasts at each time point. Values are represented as mean ± SEM. $^{**}p < 0.01$.

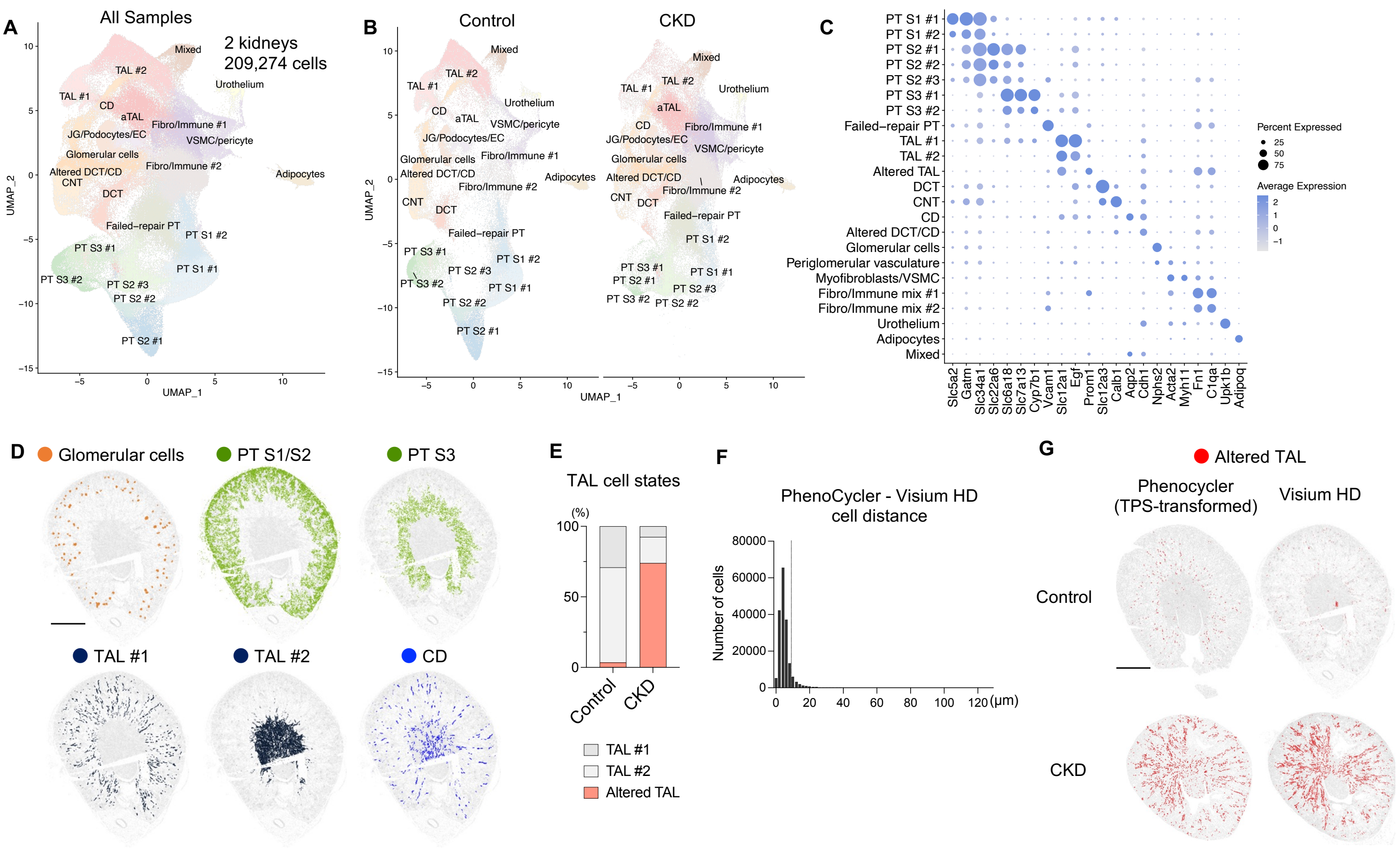


**Fig. S2** (relating to Fig. 2): Visium HD spatial transcriptome analysis was conducted on control and IRI-induced CKD kidneys. (A) UMAP plot showing cell types. (B) UMAP plot showing cell types in each condition. (C) Dot plot showing average expression of marker genes and percentage of cells expressing each marker. (D) Cellular distribution in control kidneys. Bar = 1mm. (E) Proportion of TAL subtypes in control and CKD kidneys. (F) Histogram of distances from PhenoCycler cells to their nearest Visium HD cells. (G) Spatial distribution of aTAL cells defined by PhenoCycler and by Visium HD in control and CKD kidneys. Bar = 1 mm.

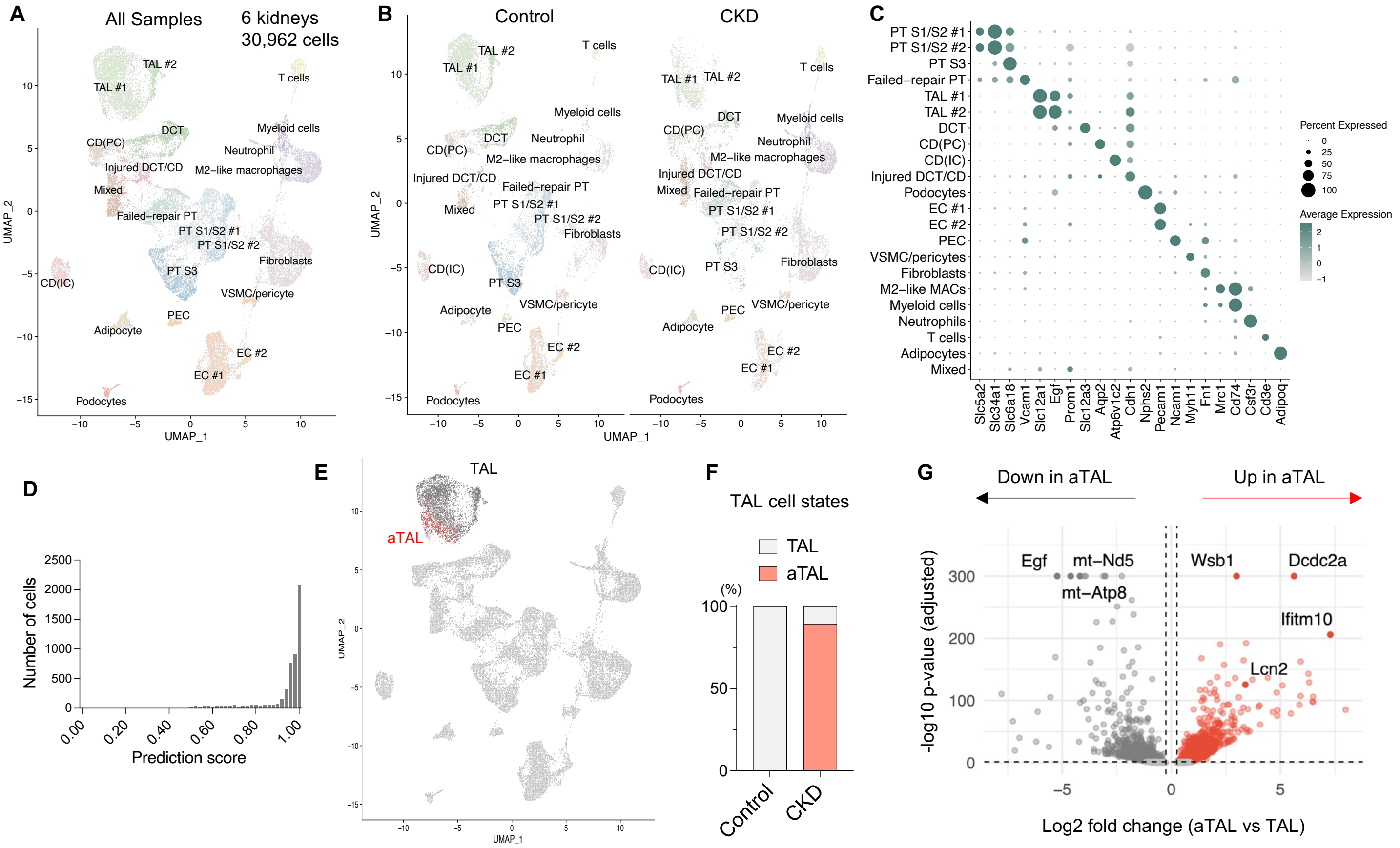


**Fig. S3** (relating to Fig. 2): GEM-X Flex Gene Expression assay was conducted on control and IRI-induced CKD kidneys. (A) UMAP plot showing cell types. (B) UMAP plot showing cell types in each condition. (C) Dot plot showing average expression of marker genes and percentage of cells expressing each marker. (D) Distribution of label transfer prediction scores for mapping Visium HD TAL subtypes onto snRNA-seq TAL cells. (E) Projection of transferred Visium HD TAL cell-type annotations onto the original snRNA-seq UMAP. (F) Proportion of transferred TAL subtypes in control and CKD kidneys. (G) Volcano plot showing differentially expressed genes in aTAL cells compared to TAL cells. Genes with adjusted $p < 0.05$ and $\log2FC > 0.25$ are shown in red, whereas genes with adjusted $p < 0.05$ and $\log2FC < -0.25$ are shown in gray. All other genes are shown in light gray. The names of the top three upregulated and top three downregulated genes, as well as *Lcn2*, are highlighted.

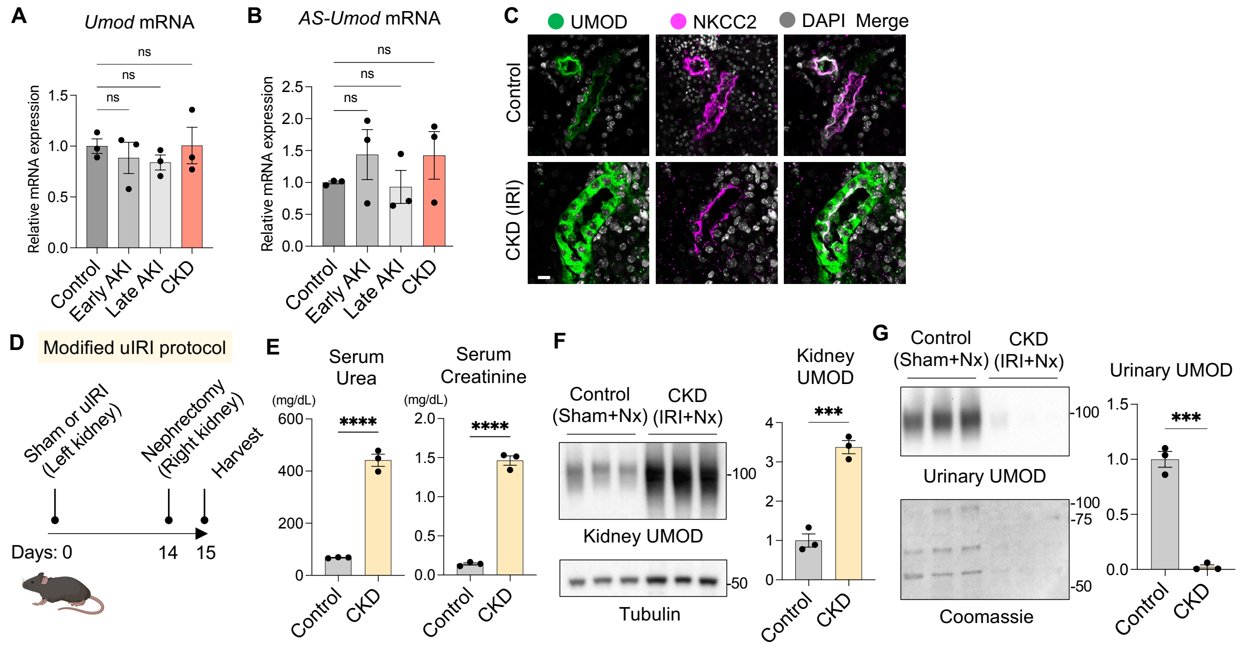


**Fig. S4** (relating to Fig. 3): (A) Relative *Umod* mRNA expression normalized to *Gapdh* in IRI kidneys. n = 3 per group. (B) Relative mRNA expression of alternatively spliced *Umod (AS-Umod)* normalized to *Gapdh* in IRI kidneys. n = 3 per group. (C) Representative immunofluorescence images of UMOD and NKCC2 in IRI kidneys. Bar = 10 µm. n = 3 per group. (D-G) To assess renal function and urinary UMOD secretion from IRI kidneys, contralateral nephrectomy (Nx) was performed 24 hours before harvest. (D) Schematic of the modified IRI experimental design. (E) Serum creatinine and urea concentration. n = 3 per group. (F) Immunoblotting of kidney UMOD. n = 3 per group. (G) Immunoblotting of urinary UMOD. The same volume of urine was applied. n = 3 per group. Values are represented as mean ± SEM. ***$p < 0.001$; ****$p < 0.0001$.

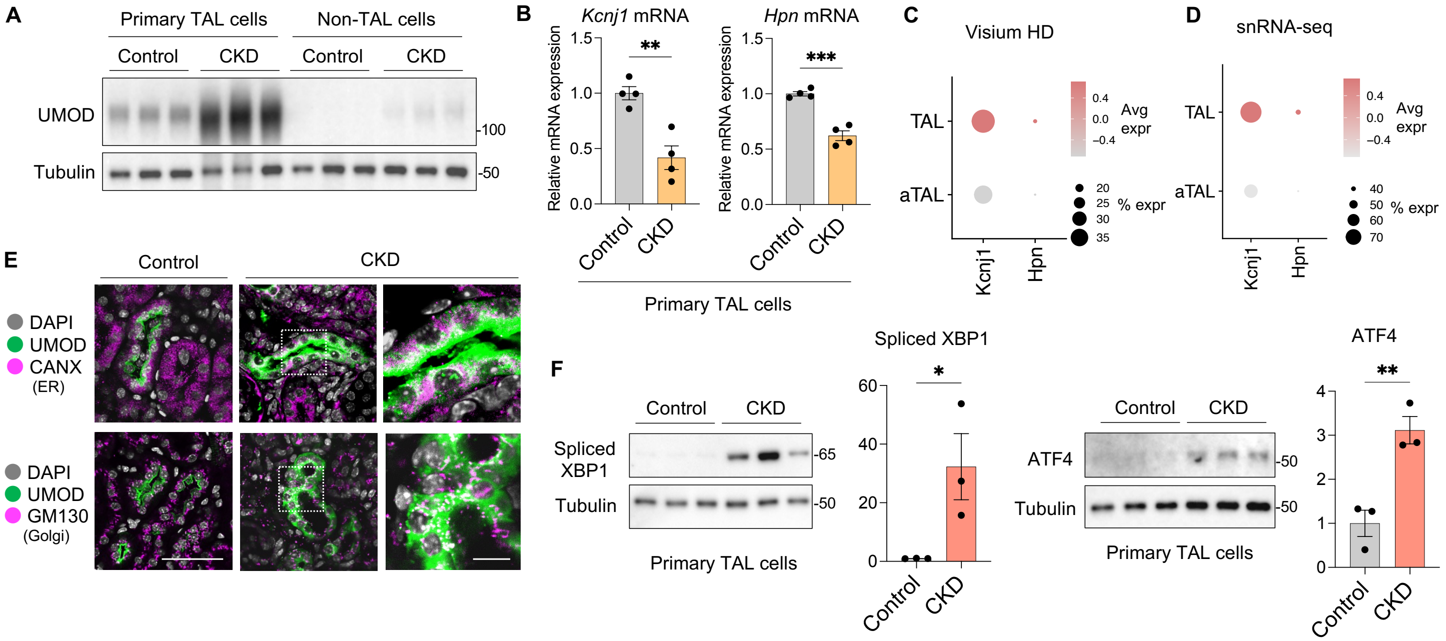


**Fig. S5** (relating to Fig. 3): (A,B) Primary TAL cells were isolated from control and CKD (IRI day 14) mouse kidneys. (A) Immunoblotting of UMOD in TAL cells and non-TAL cells. n = 3 per group. (B) qPCR analysis of *Kcnj1* and *Hpn* normalized to *Gapdh* in primary TAL cells. n = 4 per group. (C) Dot plots showing *Kcnj1* and *Hpn* expression in aTAL cells from spatial transcriptomics dataset. (D) Dot plot showing *Kcnj1* and *Hpn* expression in aTAL cells from snRNA-seq dataset. (E) Immunofluorescence images of UMOD, CANX (ER marker) and GM130 (Golgi marker). Bars = 50 μm or 10 μm (zoom in). n = 3 per group. (F) Immunoblotting of spliced XBP1 (XBP1s) and ATF4 in TAL cells in CKD. n = 3 per group. Values are represented as mean ± SEM. *p<0.05, **p<0.01, ***p < 0.001.

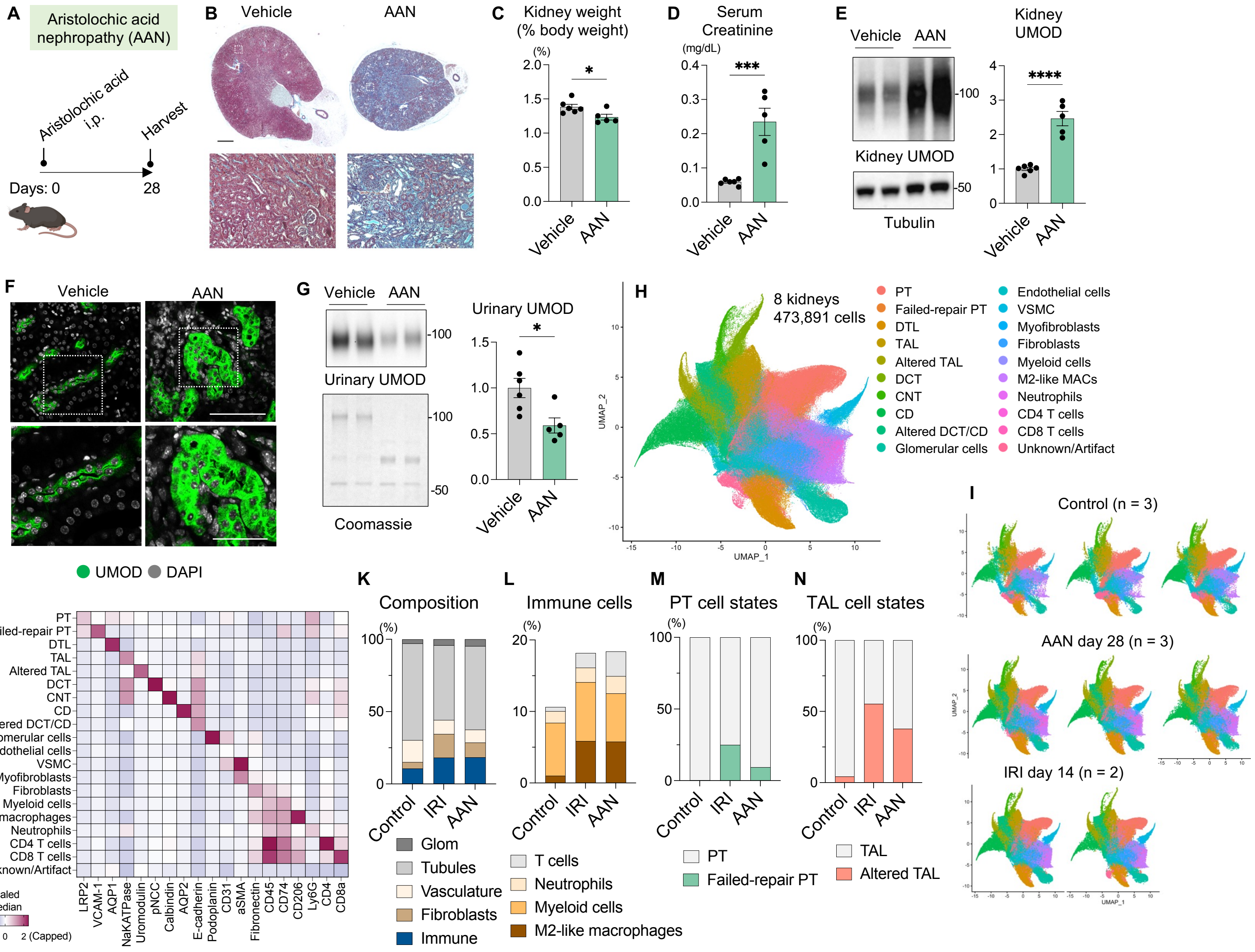


**Fig. S6** (relating to Fig. 3): (A-G) To establish aristolochic acid nephropathy (AAN) model, wild-type mice underwent a single intraperitoneal injection of aristolochic acid (5mg/kg) and were harvested 28 days after injection. n = 5-6 per group. (A) Schematic of the experimental design. (B) Masson's trichrome staining of AAN kidneys. Scale bars = 1mm and 100 μm (zoom in). n = 5-6 per group. (C) Total kidney weight (left and right) normalized to basal body weight in AAN mice. n = 5-6 per group. (D) Serum creatinine concentration. n = 5-6 per group. (E) Immunoblotting of kidney UMOD in AAN model. n = 5-6 per group. (F) Representative immunofluorescence images of UMOD in AAN kidneys. Scale bars = 10 μm and 5 μm (zoom in). n = 3 per group. (G) Immunoblotting of urinary UMOD. The same volume of urine was applied. n = 5 per group. (H-N) PhenoCycler protein imaging was performed on kidneys from uninjured control mice, IRI mice at day 14, and AAN mice at day 28 (n = 3, 2, and 3, respectively). (H) UMAP plot showing cell types. (I) UMAP plots of all individual samples. (J) Heatmap displaying median expression of marker proteins across clusters. (K) Cellular composition in each condition. (L) Proportion of immune cell subtypes in each condition. (M) Proportion of failed-repair PT cells among total PT cells in each condition. (N) Proportion of altered TAL cells among total TAL cells in each condition. Values are represented as mean ± SEM. *$p<0.05$, ***$p < 0.001$; ****$p < 0.0001$.

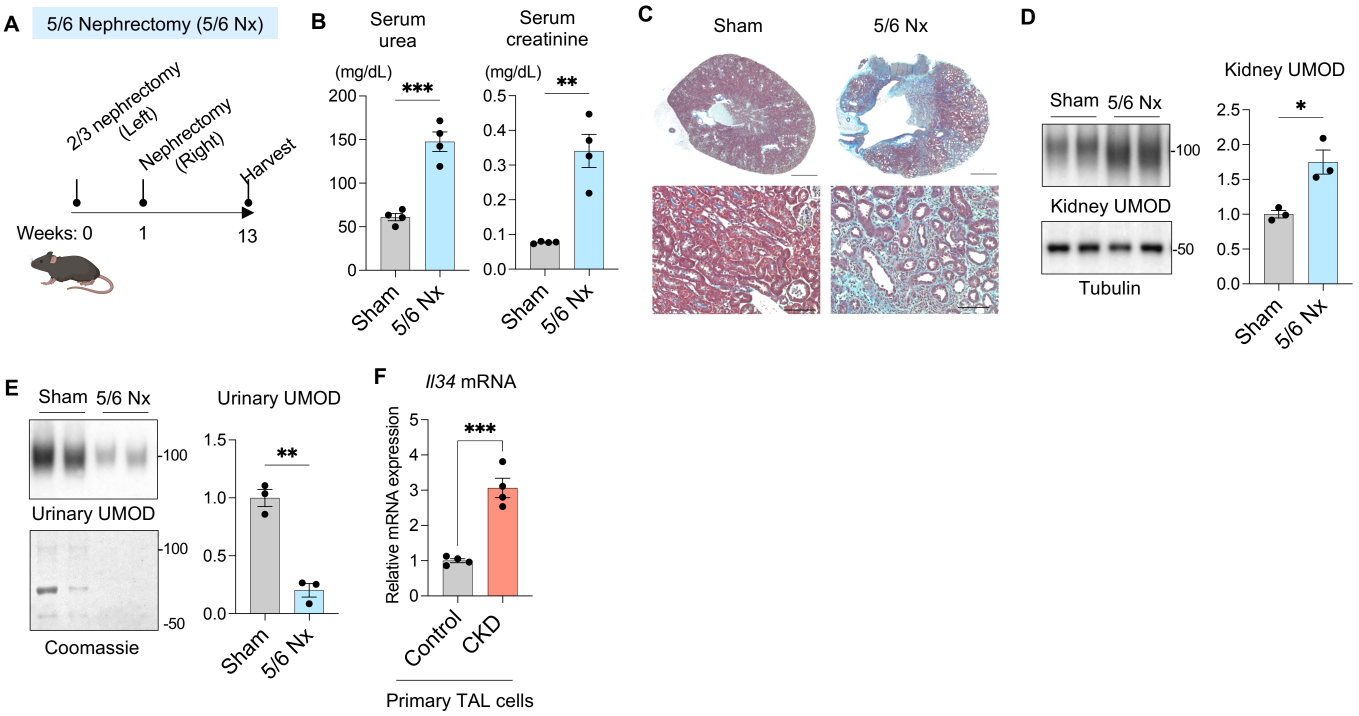


**Fig. S7** (relating to Fig. 3,4): (A-E) To establish a 5/6 nephrectomy (5/6 Nx) model, two-thirds of the left kidney was resected, followed by right nephrectomy 1 week later. Mice were harvested 12 weeks after the second surgery. (A) Schematic of the experimental design. (B) Serum urea and creatinine levels. n = 4 per group. (C) Masson's trichrome staining of 5/6 Nx kidneys. Bars = 1mm and 100 µm (zoom in). n = 4 per group. (D) Immunoblotting of kidney UMOD. n = 3 per group. (E) Immunoblotting of urinary UMOD. The same volume of urine was applied. n = 3 per group. (F) qPCR of Il34 in primary TAL cells from IRI-induced CKD kidneys. n = 4 per group. Values are represented as mean ± SEM. *p < 0.05, **p < 0.01, ***p < 0.001.

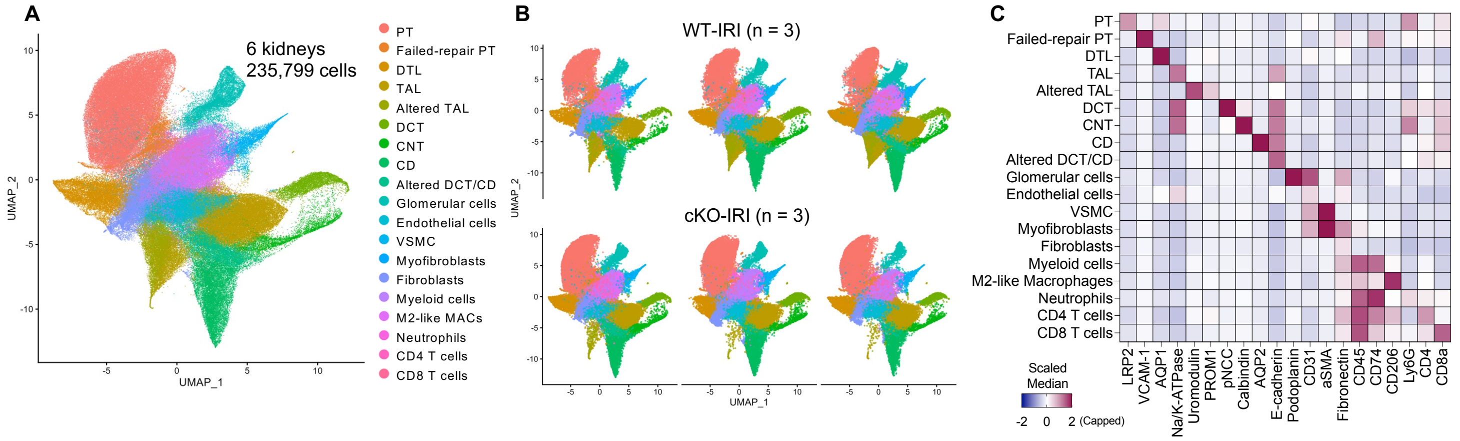


**Fig. S8** (relating to Fig. 5): (A) UMAP plot showing cell types. (B) UMAP plots of all individual samples. (C) Heatmap displaying median expression of marker proteins across clusters.

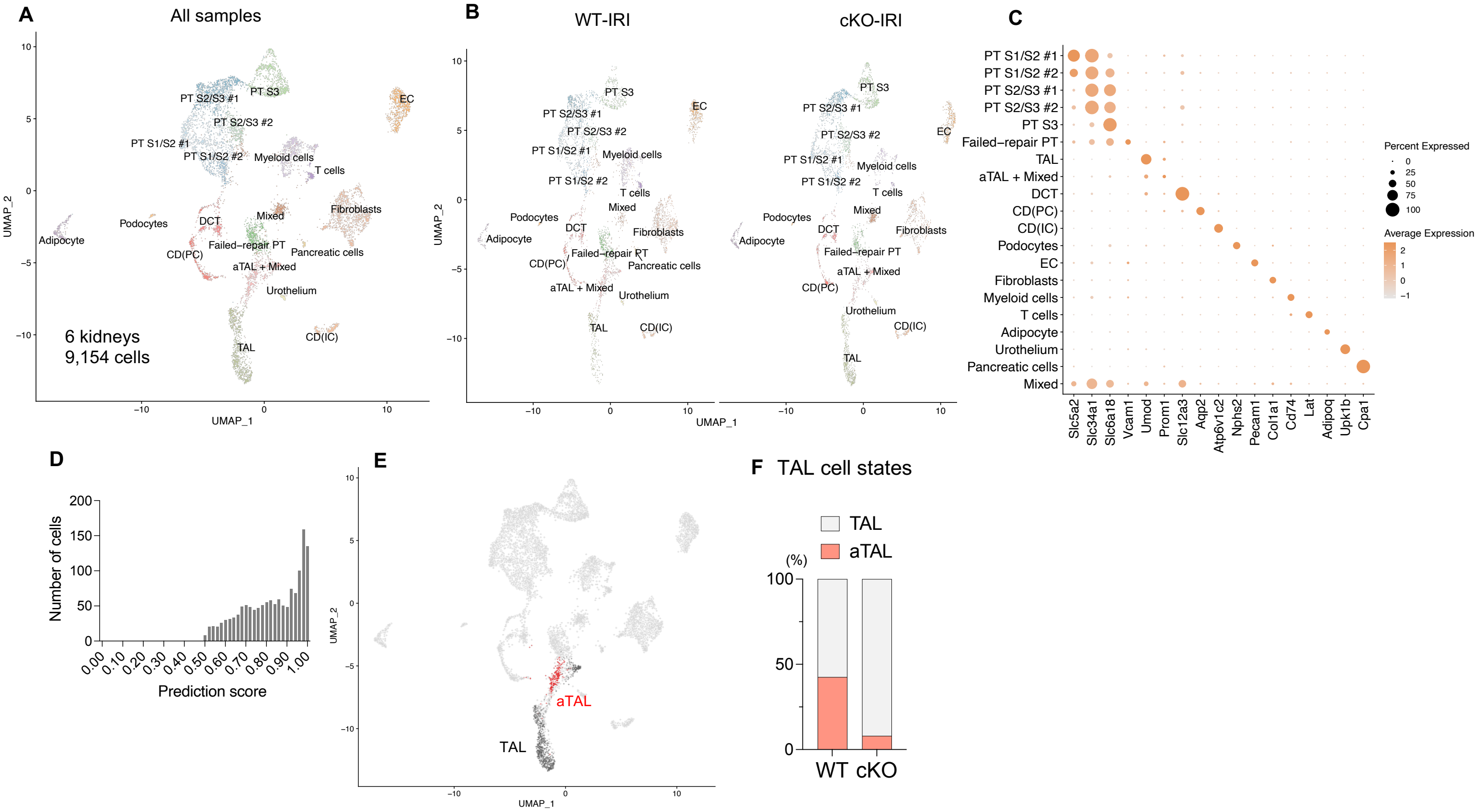


**Fig. S9** (relating to Fig. 5): GEM-X Flex Gene Expression assay was conducted on WT-IRI and cKO-IRI kidneys. (A) UMAP plot showing cell types. (B) UMAP plot showing cell types in each condition. (C) Dot plot showing average expression of marker genes and percentage of cells expressing each marker. (D) Distribution of label transfer prediction scores for mapping Visium HD TAL subtypes onto snRNA-seq TAL cells. (E) Projection of transferred Visium HD TAL cell-type annotations onto the original snRNA-seq UMAP. (F) Proportion of transferred TAL subtypes in WT-IRI and cKO-IRI kidneys.

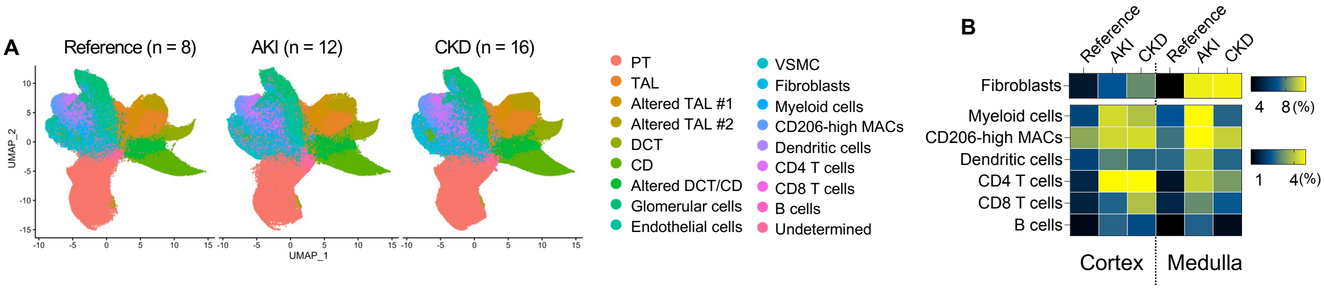


**Fig. S10** (relating to Fig. 6): (A) UMAP plots per condition. (B) Heatmap showing proportion of each cell type across conditions and anatomical regions.

| Marker | Vendor | Catalog # | Clone |
|---|---|---|---|
| AQP1 | Abcam | ab178352 | EPR11588(B) |
| AQP2 | Abcam | ab230170 | EPR21080 |
| aSMA | Invitrogen | 14-9760-82 | 1A4 |
| b-catenin | NovusBio | NBP2-80596 | 12F7 |
| Calbindin | Abcam | ab82812 | CB-955 |
| CD11c | NovusBio | NB110-97871 | AP-MAB0806 |
| CD153 | Invitrogen | 14-1531-85 | RM153 |
| CD16 | BioLegend | 302049 | 3G8 |
| CD206 | R&D Systems | AF2534 | Polyclonal (Goat) |
| CD279 (PD-1) | Invitrogen | 14-9985-82 | J43 |
| CD3 | Invitrogen | 14-0032-82 | 17A2 |
| CD31 | Abcam | ab225883 | EPR17259 |
| CD4 | Biolegend | 100401 | GK1.5 |
| CD45 | R&D Systems | AF114 | Polyclonal (Goat) |
| CD45r/B220 | Invitrogen | 14-0452-82 | RA3-6B2 |
| CD63 | Biolegend | 353039 | H5C6 |
| CD74 | R&D Systems | AF7478 | Polyclonal (Sheep) |
| CD8a | Invitrogen | MA5-13473 | C8/144B |
| Cystatin C | Abcam | Ab217569 | EPR4413 |
| E-cadherin | AbboMax | 605-730 | 4A2C7 |
| Fibronectin | Abcam | ab268023 | EPR23110-25 |
| GATA3 | Cell Signaling Technology | 10630SF | D13C9 |
| Ki67 | BD Biosciences | 556003 | B56 |
| LRP2 | Abcam | ab76969 | Polyclonal (Rabbit) |
| Ly6C | Invitrogen | 14-5931-82 | RB6-8C5 |
| Ly6G | Invitrogen | 16-9668-82 | 1A8-ly6g |
| Na/K-ATPase | GeneTex | GTX635461 | HL114 |
| Nephrin | Abcam | ab227806 | EPR20993 |
| NK1.1 | BioLegend | 108701 | PK136 |
| O-GlcNAc | NovusBio | NB300-524 | RL2 |

| | | | |
|---|---|---|---|
| PDK4 | Invitrogen | PA5-13776 | Polyclonal (Rabbit) |
| PLSCR1 | Invitrogen | MA5-19636 | 3B4 |
| pMLKL | Cell Signaling Technology | 91689BF | D6H3V |
| pNCC | Phosphosolutions | p1311-53 | Polyclonal (Rabbit) |
| Podoplanin | Invitrogen | 14-5381-81 | eBio8.1.1 |
| PROM1 | BioLegend | 394002 | S16016B(AC133) |
| S100A4 | LSBio | LS-B11817 | Polyclonal (Rabbit) |
| Sox9 | Abcam | ab220812 | EPR14335 |
| Uromodulin | R&D Systems | AF5144 | Polyclonal (Sheep) |
| VCAM-1 | Abcam | ab271899 | EPR5047 |
| VWF | NovusBio | NBP3-21110 | VWF/7979R |
| Vimentin | BD Biosciences | 550513 | RV202 |

**Table S1**: PhenoCycler antibody panel for IRI trajectory dataset.

| Marker | Vendor | Catalog # | Clone |
|---|---|---|---|
| AQP1 | Abcam | ab178352 | EPR11588(B) |
| AQP2 | Abcam | ab230170 | EPR21080 |
| aSMA | Invitrogen | 14-9760-82 | 1A4 |
| b-catenin | NovusBio | NBP2-80596 | 12F7 |
| Calbindin | Abcam | ab82812 | CB-955 |
| CD11c | NovusBio | NB110-97871 | AP-MAB0806 |
| CD153 | Invitrogen | 14-1531-85 | RM153 |
| CD206 | R&D Systems | AF2534 | Polyclonal (Goat) |
| CD279 (PD-1) | Invitrogen | 14-9985-82 | J43 |
| CD3 | Invitrogen | 14-0032-82 | 17A2 |
| CD31 | Abcam | ab225883 | EPR17259 |
| CD4 | Abcam | ab183685 | EPR19514 |
| CD45 | R&D Systems | AF114 | Polyclonal (Goat) |
| CD45r/B220 | Invitrogen | 14-0452-82 | RA3-6B2 |
| CD63 | Biolegend | 353039 | H5C6 |
| CD74 | R&D Systems | AF7478 | Polyclonal (Sheep) |
| CD8a | Invitrogen | MA5-13473 | C8/144B |
| Collagen 1a1 | Abcam | AB215969 | EPR7785 |
| CPT1a | Proteintech | 15184-1-AP | Polyclonal (Rabbit) |
| Cystatin C | Abcam | Ab217569 | EPR4413 |
| E-cadherin | AbboMax | 605-730 | 4A2C7 |
| F4/80 | Abcam | ab300422 | EPR26545-166 |
| Fibronectin | Abcam | ab268023 | EPR23110-25 |
| GATA3 | Cell Signaling Technology | 10630SF | D13C9 |
| Ki67 | BD Biosciences | 556003 | B56 |
| LRP2 | Abcam | ab76969 | Polyclonal (Rabbit) |
| Ly6C | Invitrogen | 14-5931-82 | RB6-8C5 |
| Ly6G | Invitrogen | 16-9668-82 | 1A8-ly6g |
| MCP-1 | Santa Cruz Biotech | sc-32771 | 5J |
| Na/K-ATPase | GeneTex | GTX635461 | HL114 |

| | | | |
|---|---|---|---|
| Nephrin | Abcam | ab227806 | EPR20993 |
| O-GlcNAc | NovusBio | NB300-524 | RL2 |
| PLSCR1 | Invitrogen | MA5-19636 | 3B4 |
| pMLKL | Cell Signaling Technology | 91689BF | D6H3V |
| pNCC | Phosphosolutions | p1311-53 | Polyclonal (Rabbit) |
| Podoplanin | Invitrogen | 14-5381-81 | eBio8.1.1 |
| PROM1 | Proteintech | 18470-1-AP | Polyclonal (Rabbit) |
| S100A4 | LSBio | LS-B11817 | Polyclonal (Rabbit) |
| Sox9 | Abcam | ab220812 | EPR14335 |
| Uromodulin | R&D Systems | AF5144 | Polyclonal (Sheep) |
| VCAM-1 | Abcam | ab271899 | EPR5047 |
| VWF | NovusBio | NBP3-21110 | VWF/7979R |

**Table S2**: PhenoCycler antibody panel for IRI and AAN dataset and Umod cKO mice dataset.

| | | Reference | AKI | CKD |
|---|---|---|---|---|
| Number of individuals | | 8 | 12 | 16 |
| Number of cortex specimen | | 8 | 11 | 15 |
| Number of medullary specimen | | 2 | 2 | 7 |
| Age | | 39.0 ± 10.9 | 44.7 ± 16.0 | 61.6 ± 13.6 |
| Gender | Male | 2 | 12 | 6 |
| | Female | 6 | 0 | 10 |
| Race | Asian | 1 | 0 | 0 |
| | Black or African-American | 0 | 3 | 5 |
| | Other | 0 | 1 | 0 |
| | White | 7 | 8 | 11 |
| Comorbidities | Diabetes | 0 | 4 | 14 |
| | Hypertension | 0 | 5 | 15 |

**Table S3**: Demographics of reference and disease kidney tissue specimens. Age was presented as mean ± SD.

**Data S1 (separate file)** : List of genes upregulated in aTAL cells.
**Data S2 (separate file)** : List of genes downregulated in aTAL cells.
